# The Impact of Gulf Stream Frontal Eddies on Ecology and Biogeochemistry near Cape Hatteras

**DOI:** 10.1101/2023.02.22.529409

**Authors:** Patrick Clifton Gray, Jessica Gronniger, Ivan Sayvelev, Julian Dale, Alexandria K. Niebergall, Nicolas Cassar, Anna E. Windle, Dana E. Hunt, Zackary Johnson, Marina Lévy, Chris Taylor, Guillaume Bourdin, Ashley Blawas, Amanda Lohmann, Greg Silsbe, David W. Johnston

**Affiliations:** Duke University Marine Laboratory, Nicholas School of the Environment, Duke University, Beaufort NC, USA; School of Marine Sciences, University of Maine, Orono, ME, USA; U.S. Naval Research Laboratory, Washington, District of Columbia, USA; Nicholas School of the Environment, Duke University, Durham NC, USA; Horn Point Laboratory, University of Maryland Center for Environmental Science, Cambridge, MD, USA; Laboratoire d’Océanographie et du Climat (LOCEAN), Institut Pierre, Sorbonne Université (CNRS/IRD/MNHN), Simon Laplace (IPSL), Paris, France; NOAA National Centers for Coastal Ocean Science, Beaufort, NC, USA

## Abstract

Ocean physics and biology can interact in myriad and complex ways. Eddies, features found at many scales in the ocean, can drive substantial changes in physical and biogeochemical fields with major implications for marine ecosystems. Mesoscale eddies are challenging to model and difficult to observe synoptically at sea due to their fine-scale variability yet broad extent. In this work we observed a frontal eddy just north of Cape Hatteras via an intensive hydrographic, biogeochemical, and optical sampling campaign. Frontal eddies occur in western boundary currents around the globe and there are major gaps in our understanding of their ecosystem impacts. In the Gulf Stream, frontal eddies have been studied in the South Atlantic Bight, where they are generally assumed to shear apart passing Cape Hatteras. However, we found that the observed frontal eddy had different physical properties and phytoplankton community composition from adjacent water masses, in addition to continued cyclonic rotation. In this work we first synthesize the overall ecological impacts of frontal eddies in a simple conceptual model. This conceptual model led to the hypothesis that frontal eddies could be well timed to supply zooplankton to secondary consumers off Cape Hatteras where there is a notably high concentration and diversity of top predators. Towards testing this hypothesis and our conceptual model we report on the biogeochemical state of this particular eddy connecting physical and biological dynamics, analyze how it differs from Gulf Stream and shelf waters even in “death”, and refine our initial model with this new data.

**Key Points:**

- In-depth investigation of a frontal eddy in the Gulf Stream off Cape Hatteras, North Carolina
- Continued physical and biogeochemical differences are observed between the eddy and adjacent water masses even as it begins to shear apart
- We share a conceptual model of the ecological impact of frontal eddies with a hypothesis that they supply zooplankton to secondary consumers

**Plain Language Summary:** Frontal eddies are spinning masses of water (~30km in diameter) that move along western boundary currents like the Gulf Stream. When they form they carry productive coastal water into the Gulf Stream and drive upwelling within their cores. Together this leads to an increase in the amount of phytoplankton within them - much higher compared to surrounding nutrient-limited Gulf Stream water. On the east coast of the United States one common area of frontal eddy formation is just off Charleston, SC. Eddies then travel up the coast and dissipate near Cape Hatteras, NC. In this work we measured a wide range of physical and biological properties of a frontal eddy just north of Cape Hatteras. We compared these properties within the eddy to the coastal water on one side and the Gulf Stream water on the other, finding clear differences in phytoplankton community composition and other physical and chemical properties. Using the results of these observations together with previous studies we share a simple model for how frontal eddies may impact phytoplankton, zooplankton, and fish – hypothesizing that they may contribute to the high diversity and density of top predators off Cape Hatteras.

## 1 Introduction

Circular currents of water, termed eddies, can trap and transport water properties throughout the ocean, containing and moving hydrographic and biogeochemical properties laterally as they are advected by major currents and vertically by raising or lowering density surfaces (McGillicuddy, 2016). Mesoscale eddies, with spatial scales O(100km) and temporal scales O(months), are often considered the “weather of the ocean” and are a discrete example of biophysical interaction where physics can strongly influence ocean ecology (Clayton et al., 2013; Gaube et al., 2014; Lévy, 2008; Mahadevan, 2016; Williams & Follows, 1998).

The physical processes through which eddies impact ocean ecosystems include trapping and lateral advection, stirring, upwelling and downwelling, and stratification though the ecological impacts of these processes can be complex and contradictory, for a comprehensive review see McGillicuddy (2016). As a specific example, cyclonic eddies from western boundary systems (in the Gulf Stream often called cold core rings) typically have enhanced chlorophyll *a* (chl-a) both from entrainment of more nutrient rich and productive coastal water and enhanced upwelling due to isopycnal uplift (Gaube et al., 2014), but with strong winds this could be balanced or even trend negative by wind/eddy Ekman pumping.

Given the range of physical and nutrient changes due to eddies, we expect an even more heterogeneous impact on primary productivity, phytoplankton community composition, and even higher trophic levels. Generally where the ecosystem is nutrient limited, cyclonic eddies are associated with increases in phytoplankton size, diversity, and productivity, which lead to increases in zooplankton populations (Belkin et al., 2022; Landry et al., 2008). For long-lived eddies (>6 months), modeling work suggests reduced phytoplankton diversity on average due to competitive exclusion, though individual eddies were widely variable (Lévy et al., 2015). Little work has been done on secondary consumer responses to eddies, though it is likely that after an eddy-induced bloom, transfer efficiency increases as the number of trophic links between the primary consumers and forage fish decreases, e.g. larger phytoplankton such as diatoms consumed by large zooplankton who are consumed directly by fish (Eddy et al., 2021). Eddy-driven productivity and biophysical changes can move up the trophic ladder, structuring the distribution of top predators both due to physiological preferences - anticyclonic eddies deepen the warmer mixed layer allowing sharks to access deeper prey with less temperature stress (Braun et al., 2019; Gaube et al., 2018) - and increased prey density. Thus, an increase in foraging efficiency may explain preferences for eddies in tuna, swordfish, and seabirds (Arostegui et al., 2022; Haney, 1986; Hsu et al., 2015).

One class of eddies that are relatively underexplored are frontal eddies, cyclonic features that form in the trough of meanders in western boundary currents such as the Gulf Stream, the Loop Current in the Gulf of Mexico (Maul et al., 1974; Rudnick et al., 2015), the Kuroshio Current (Kasai et al., 2002), and the East Australian Current (Ribbe et al., 2018). On the U.S. eastern seaboard, these features are important for production on the outer shelf of the South Atlantic Bight (T. N. Lee et al., 1991), but they are typically thought to shear apart when the Gulf Stream narrows and rounds Cape Hatteras, North Carolina shortly before the current separates from the continental shelf (Figure 1). In this work we investigated a frontal eddy northeast and downstream of Cape Hatteras, at the Gulf Stream’s separation point from the continental shelf. Our goals in this work were to 1) better understand the evolution of primary producers and primary consumers within frontal eddies and 2) re-examine the overall ecosystem impact of frontal eddies throughout their lifetime.

**Figure 1.**
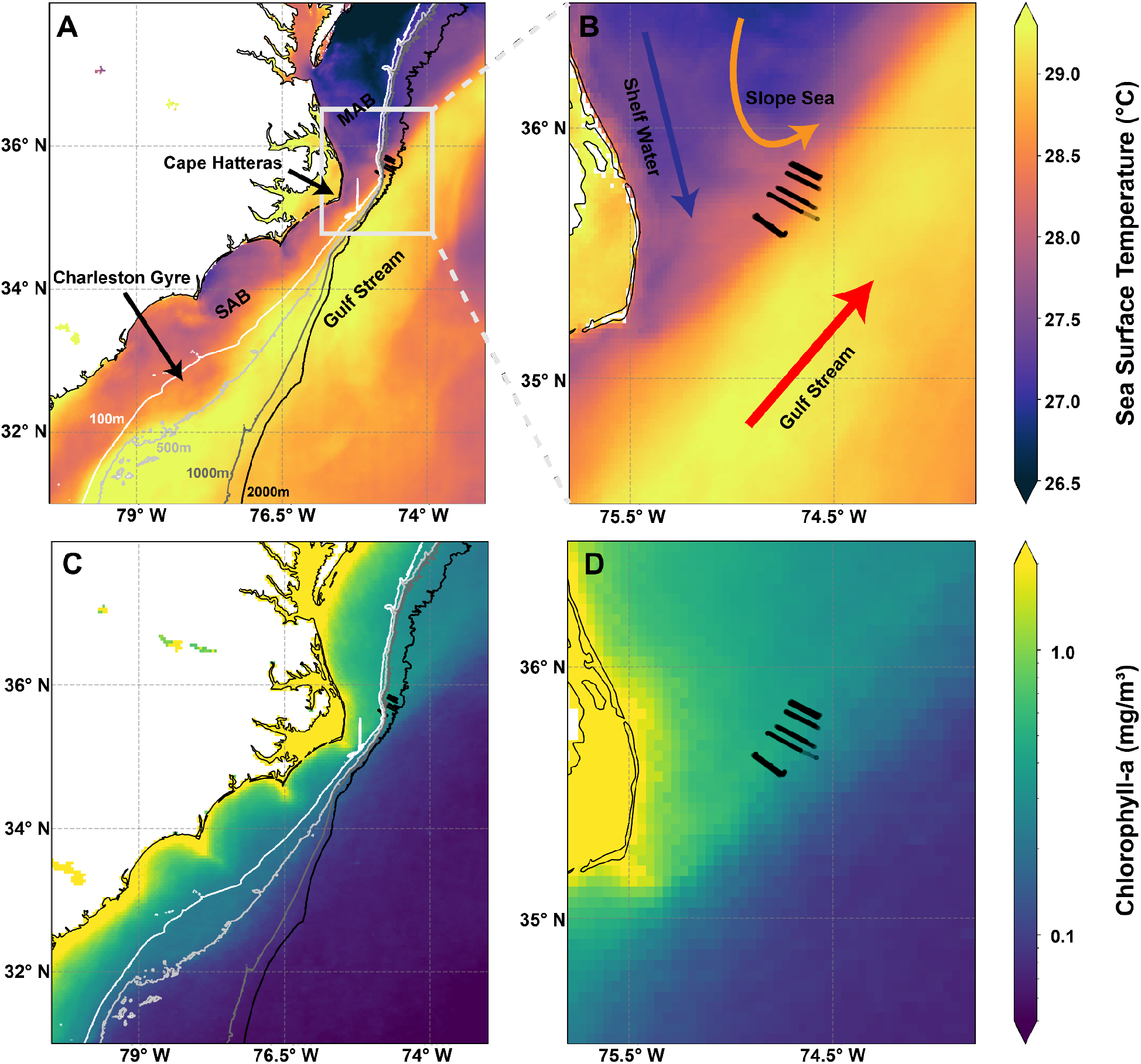
Study area overview. Panel A shows the broad geographic context using an average sea surface temperature and chlorophyll-a from August 20th to September 10th, the approximate lifetime of our eddy. Bathymetry is shown via the 100m, 500m, 1000m, and 2000m contours. The Gulf Stream is visible as the highest temperature water approximately following the 500m contour until Cape Hatteras. The Charleston Bump is visible in the 500m contour at approximately (34° N, 79° W) and the seaward deflection of the Gulf Stream is visible as a decrease in SST just north of this point. Panel B shows an inset zoom in of our study region with the nine transects shown in black. Panels C and D show the average chlorophyll-a for the same period. SST is from GOES-16 and chl-a is from the Ocean Colour Climate Change Initiative’s multi-sensor global satellite chlorophyll-a product.

Our in-depth in-situ investigation takes place around “the Point” off Cape Hatteras (Figure 1) using a combination of physical, molecular, chemical, and optical methods to elucidate the effects of a cyclonic eddy on ocean biogeochemistry. Given that the in-situ component takes place over just three days - viewed in this work as effectively a single time step - we use satellite data to complement this in-situ view and investigate the two week-long life of this eddy.

### 1.1 A Brief History of Frontal Eddy Research

There is a long, though intermittent, history of studying frontal eddies in the Gulf Stream (von Arx et al., 1955; Pillsbury, 1890; Webster, 1961). Extensive physical surveys describe these features as cyclonic, cold-core eddies with substantial upwelling through isopycnal uplift (Bane et al., 1981; Thomas N. Lee et al., 1981). In turn, phytoplankton respond intensely to this nutrient upwelling, with observed levels of diatom blooms and chl-a 10-100 times greater than those typically measured in Gulf Stream or outer shelf water (defined as depths from 40-200m (T. N. Lee et al., 1991)). Menhaden and bluefish migrations to the South Atlantic Bight (SAB) are suggested to be secondary to this response of primary producers as the high-levels of phytoplankton provide a consistent food source for higher-level consumers (Yoder et al., 1981). The Frontal Eddy Dynamics experiment in 1987 intensively surveyed a frontal eddy between Cape Lookout and Cape Hatteras, providing some of the first evidence that these features propagate northward and downstream beyond Cape Hatteras, with cross shore and along shore temperature profiles demonstrating the extent of isotherm doming and continued upwelling (Glenn & Ebbesmeyer, 1994b). This work suggested that not only did the feature move past Cape Hatteras, but upwelling continued and a second warm filament formed beyond the cape. The greatest temperature anomalies from upwelling were measured to occur around ~150m depth. Later, a longer survey of frontal eddies in the region confirmed the movement of these eddies north of Cape Hatteras, typically one every 3-7 days, and suggested they are an important and frequent mechanism for transfer from the SAB and Gulf Stream into the Mid Atlantic Bight (MAB) slope sea (Glenn & Ebbesmeyer, 1994a). More recent work has demonstrated that Gulf Stream meanders (including frontal eddies) from the SAB propagate past Cape Hatteras and slowly decay on their way from the Charleston Bump to The Point (Andres, 2021). Yet beyond this work and a few others (Churchill & Cornillon, 1991), the literature often views the North Carolina coast from Cape Lookout northward as a frontal eddy graveyard and thus overlooks the potential for ongoing ecological impacts of these features.

Frontal eddies form where energy is transferred from the mean flow of the current to eddy kinetic energy due to instability processes, often influenced by the local bathymetry. In the northern SAB (Charleston to Cape Lookout, where the eddy in this study formed), this energy transfer is enhanced by the Charleston Bump which deflects the Gulf Stream offshore (Figure 1). When small Gulf Stream meanders with positive vorticity pass through this region, this energy transfer forms mesoscale frontal eddies (see Gula et al 2015 for in depth dynamics). While typically classified as mesoscale eddies, frontal eddies differ from classic Gulf Stream rings as they do not detach from the Gulf Stream, but instead remain trapped within the meander of the current. The main body of the eddy is coastal water entrained by the meander with an upwelled cold core due to the cyclonic rotation. A shallow warm streamer that flows from the downstream meander crest upstream and around this entrained coastal water separates the eddy from other coastal water (Thomas N. Lee et al., 1981). In other regions, specific topographic and current patterns drive energy transfer but the general mechanism and result are similar e.g. East Australian Current (Schaeffer et al., 2017), Kurioshio Current (Kimura et al., 1997).

Because they are trapped, frontal eddies are subject to shear between Gulf Stream, shelf waters, and dramatic bathymetry gradients. Frontal eddies are also much less nonlinear than typical mesoscale eddies (defined as U/c, the ratio of rotational speed U to the translational speed c of the feature). When U/c > 1 the feature is defined as nonlinear, indicating increased coherence. While a typical Gulf Stream ring might be 0.2/0.02 m/s ~10, a frontal eddy could be approximately 0.5/0.5 m/s ~1 (Glenn and Ebbesmeyer 1994a), suggesting it is less likely to keep a coherent structure as it propagates downstream (Chelton et al., 2011). Differences between frontal eddies and Gulf Stream rings are also described by differences in their Rossby number (Ro = U/*f*L, where U – characteristic velocity, L – characteristic length scale, *f* – Coriolis frequency), where a smaller Ro indicates a more stable eddy (dominated by geostrophic balance). For our frontal eddy Ro = 0.5 / (2.5e-5*3e4) = 0.66 compared to a typical Gulf Stream ring at the same latitude where Ro = 0.2 / (2.5e-5*1e5) = 0.008 suggesting less stability. Rossby numbers for other frontal eddies such as those in the East Australian Current have been reported to be similar or higher (0.6-1.9) (Schaeffer et al., 2017). Compared to other mesoscale eddies, frontal eddies are typically smaller (20-50 km vs. 100+ km), shorter lived (2-3 weeks vs. 6-18 months), and form all year at a higher rate (once every 3-7 days). They often occupy nearly half or even a majority of the Gulf Stream-shelf edge in the SAB, thus forming an effective and possibly dominant mechanism for exchange between the Gulf Stream and the shelf (Gula et al., 2015).

Depths of frontal eddies are typically 50-200m (Thomas N. Lee, 1975) along stream lengths that are 2-3x cross stream. Frontal eddies in this area typically travel at the same speed as the Gulf Stream meander and are thought to have upwelling on the order of 10 m/day within 200m depth. Important to this upwelling is the deep nutrient core of the Gulf Stream, found below 100-200m, with concentrations of nitrate >10um (Pelegri & Csanady, 1991). Frontal eddies pump the nutrient core up into the euphotic zone via isopycnal uplift driving important nutrient fluxes into the euphotic zone. For eddies formed around the Charleston Bump, this generates considerable productivity from Charleston to Cape Hatteras (T. N. Lee et al., 1991). Recent high resolution modeling (*d*x=150m) shows frontal eddies contain a rich submesoscale field, localized upwelling, and numerous submesoscale eddies forming on the edge of the frontal eddy and the Gulf Stream (Gula et al., 2016) with the possibility for major additional biological implications (Lévy et al., 2018).

While the physical and hydrographic dynamics of frontal eddies are reasonably well understood, how these dynamics impact ecosystem function and composition is not well known. Most work concludes that these eddies are sheared apart from the Gulf Stream and left stranded on the outer shelf of the SAB (Lee 1981, 1991), but our work and a few previous studies show that some of these eddies, though they do experience shear from Cape Lookout to Cape Hatteras, are still partially coherent and transport their contents to the MAB and possibly far downstream. In fact, the satellite record shows that a large number of these eddies maintain some coherence past Cape Hatteras and the remnants of frontal eddies and their warm streamers can often be seen in SST imagery as far as the Scotian Shelf (Glenn & Ebbesmeyer, 1994a). Some limited work into the ecosystem implications of frontal eddies in the SAB indicates that the upwelled nutrients are consumed within 2 weeks and this may drive a zooplankton bloom (McClain & Atkinson, 1985; Paffenhöfer et al., 1987), though that work assumes nitrate was primarily being fluxed onto the shelf and the eddy was fully dissipating.

Based on the literature we developed a simple qualitative conceptual model as a framework for our study (Figure 3). In this model we expect to see a bloom of phytoplankton soon after eddy formation, followed by a zooplankton bloom around a week after the phytoplankton peak, and increased secondary consumers soon after the zooplankton peak. We expect the biology to lag the physics such that phytoplankton growth above the baseline follows upwelling by a day or two and terminates soon after upwelling ends. Our study investigates a single point in time towards the end of the eddy lifetime. Specifically, we examine how physical and biological properties are spatially structured and differ across the Gulf Stream, frontal eddy, MAB slope, and MAB shelf waters and use this along with satellite data, and previous work to understand the natural history of Gulf Stream frontal eddies.

### 1.3 Frontal Eddies and the Gulf Stream’s separation point

The in-situ focus area for this study is the confluence of the warm and salty SAB, the relatively fresh and cool MAB slope sea water, the even fresher and cooler MAB shelf waters, and the saltiest and hottest Gulf Stream itself (Seim et al., 2022). The anticyclonic rotation of the subtropical gyre, of which the Gulf Stream is the western expression, converges here with the cyclonic rotation of the slope sea gyre, along with inputs from the MAB shelf water and SAB shelf water (which typically converge at the Hatteras Front). It is a known biodiversity hotspot with some of the highest marine mammal diversity in the world (Byrd et al., 2014), major fisheries for snappers, groupers, tunas, and mackerels, and recreational fishing due to the high density of major sport fish such as tunas and other billfish.

We surveyed a frontal eddy just northeast of Cape Hatteras (Figures 1 and 2) in September 2021 with a comprehensive set of tools observing the physical, chemical, and biological status of the region and during both pre-eddy conditions and across the middle of the eddy.

**Figure 2.**
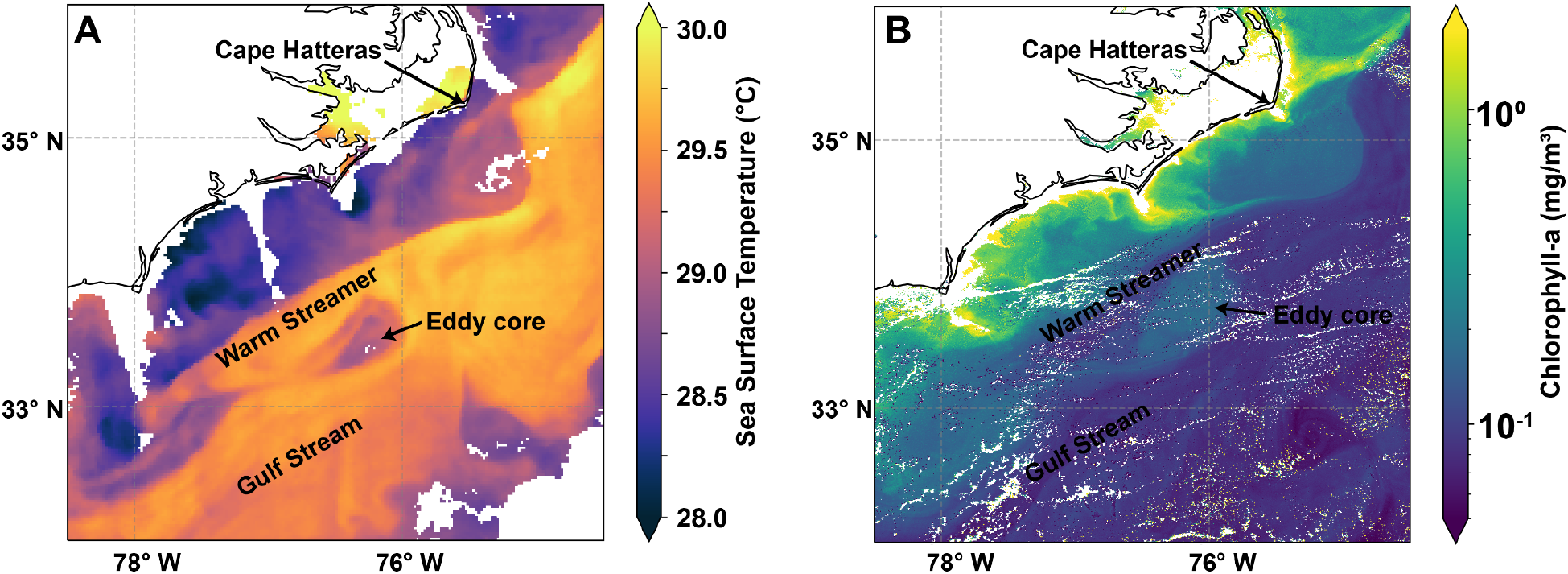
Sea surface temperature (Panel A, SST) and chlorophyll *a* (Panel B, chl-a) imagery annotated to show the structure of the sampled frontal eddy on August 31st, 2021. The warm streamer is clearly visible as is the cooler and higher chl-a eddy core. The chl-a image was acquired 13 hours after the SST image so the eddy is slightly further downstream. Another frontal eddy is visible downstream off of Cape Hatteras, though the warm streamer has been pulled into the eddy core and mixed away but is still clearly evident from both the meander in the stream, the beginning of the formation of a new warm streamer, and the positive chl-a anomaly.

**Figure 3.**
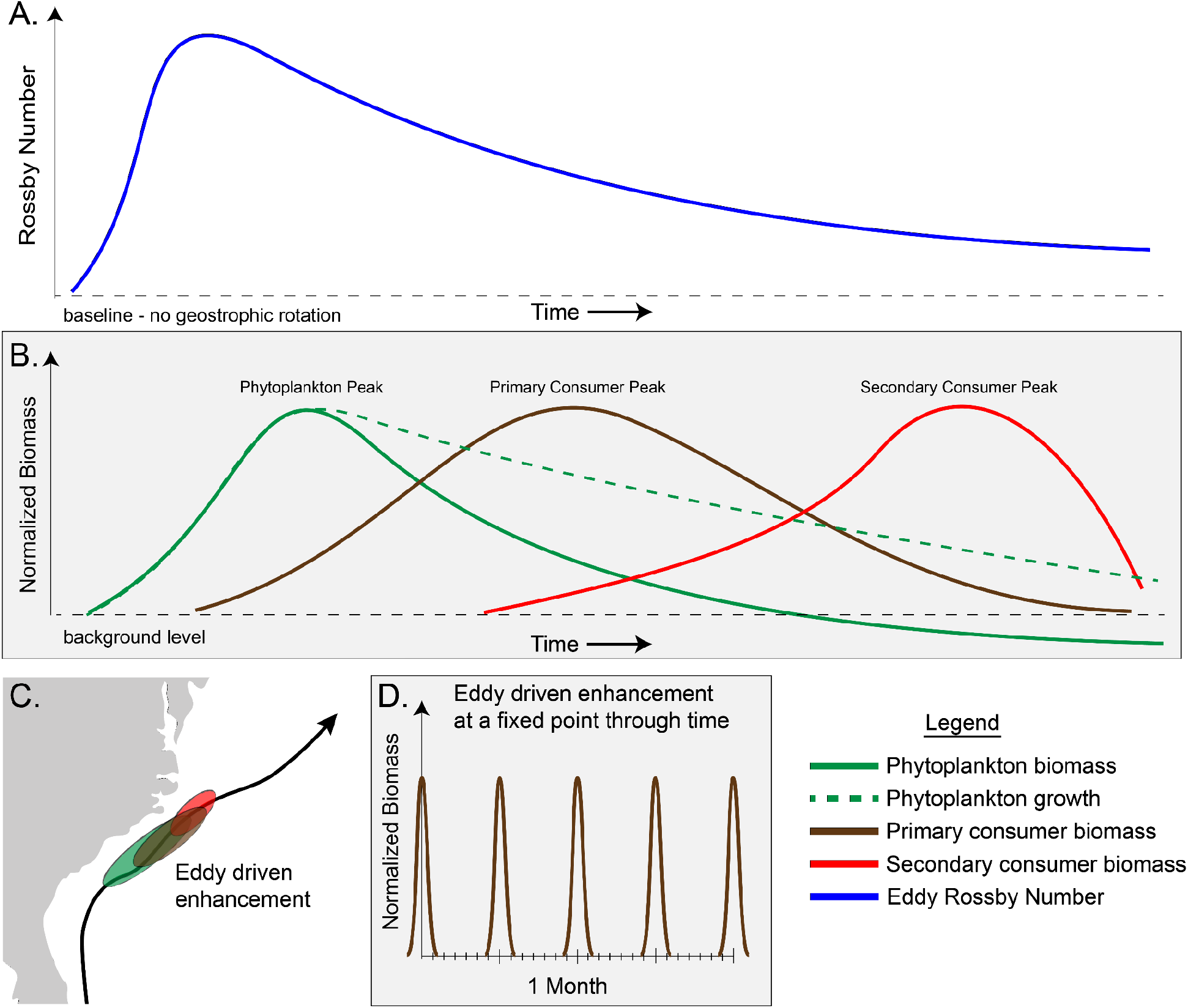
Conceptual model of a frontal eddy from the SAB to the MAB. In Panel A the eddy’s Rossby number is a proxy for geostrophic based upwelling and peaks just after formation and slowly dissipates as it moves downstream. In Panel B phytoplankton biomass peaks soon (3-5 days) after this peak in upwelling, zooplankton then peaks soon after the phytoplankton peak (5-7 days), and secondary consumer biomass in the eddy peaks a few days after zooplankton. Importantly phytoplankton growth is still likely elevated due to upwelling of nutrients even as standing biomass returns to baseline levels or even below due to grazing. Additionally while phytoplankton and zooplankton biomass is grown in-place, secondary consumer biomass is not and instead represents visits. Panel C: while the exact location of these enhancements isn’t clear and is likely variable, this approximately puts the zooplankton enhancement just off Cape Lookout and Cape Hatteras. Panel D: from a fixed perspective at this peak location, this process manifests as an enhancement of zooplankton for ~1 day of every 3-7 days year round.

The overall objective of this work is to first investigate the biogeochemical status of this frontal eddy past Cape Hatteras, where frontal eddies have rarely been sampled, particularly to see if it contains a different phytoplankton community composition than adjacent waters. Second we assess if the observations suggest an enhancement of grazers that could be supplying secondary consumers in this region. And third we use this new understanding to improve our model of the ecological evolution of eddies from formation to dissolution and their impact on the outer shelf and Gulf Stream ecosystems where they transit.

## 2 Materials and Methods

### 2.1 Focal Region

The eddy studied in this work formed off the Charleston Bump on August 25th-26th, moved past Cape Hatteras on Sept 4th, was surveyed on Sept 5th, 6th, and 7th and moved rapidly downstream and was dissipating, but still apparent by Sept 9th.

### 2.2 Ship transects

Transects on the R/V Shearwater transited the North Wall of the Gulf Stream in approx 10-15 km lines with five data intensive day time transects and four less data intensive night time transects. Transects were planned to cross the front between the shelf and eddy or shelf and Gulf Stream water. In this work we collected nine transects, five in the daytime with a more intensive approach, and four at night (Figures 1, 4 and S1).

**Figure 4.**
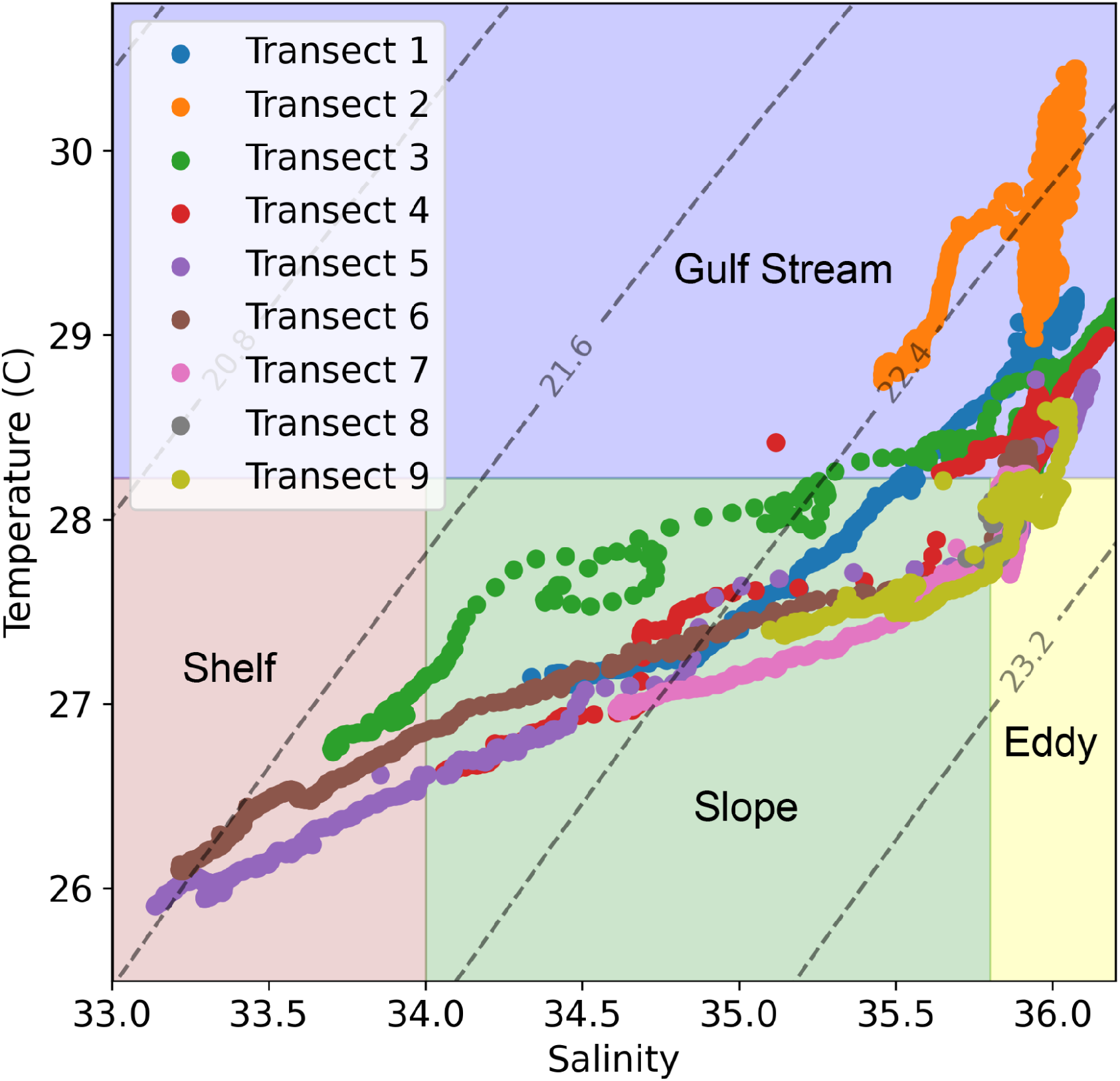
All nine transects are plotted together on a Temperature-Salinity diagram. Isopycnals are shown with dashed gray lines across the plot. The reddish section with low temperature and low salinity is the shelf water, green is the slope water, yellow is the eddy, and blue is the Gulf Stream.

### 2.3 Temperature and Salinity

Temperature and salinity were collected with a SeaBird SBE38 Thermosalinograph collected via the Shearwater’s flow through system from an intake at approximately 1.5m depth. This data was logged once per second.

### 2.4 Chl-a sampling

In-situ chl-a was measured by filtering 100 mL of seawater onto combusted 0.7 μm GF/F filters, extracting in 100% methanol for 48 hours, and reading fluorescence on a chlorophyll calibrated Turner 10AU fluorometer equipped with Welschmeyer filters (Johnson et al., 2010).

### 2.5 Flow cytometry

Duplicate whole seawater samples were collected from the ship’s flow through system at 1m depth, fixed with net 0.125% glutaraldehyde and stored at −80 °C until processing. Prokaryotic phytoplankton populations were enumerated using a Becton Dickinson FACSCalibur Flow Cytometer and categorized as previously described (Johnson et al., 2010). Bacterioplankton were quantified using SYBR Green-I on the Attune NxT acoustic flow cytometer (Life Technologies) (Marie et al., 1997).

### 2.6 Nutrients

Nutrients were collected by filtering duplicate ~50mL samples through 0.22 *μ*m Sterivex filters which was stored at −80 °C until analysis at the UCSD Nutrient Analysis facility (https://scripps.ucsd.edu/ships/shipboard-technical-support/odf/chemistry-services/nutrients) where a Seal Analytical continuous-flow AutoAnalyzer-3 was used to measure silicate, nitrate, nitrite, phosphate, and ammonia using the analytical methods described by (Atlas et al., 1971; Gordon et al., 1992; Hager et al., 1972).

### 2.7 Optical and CDOM

Hyperspectral absorption (a), attenuation (c), and the volume scattering function (VSF) at 120 deg and 470, 532, and 650 nm were measured continuously (4 Hz, 4 Hz, and 1 Hz respectively) for the duration of this work. a and c were measured at 81 wavelengths 399 to 736 nm using a WetLabs ACS spectrophotometer and VSF was measured at 120 deg at 470, 532, and 650 nm using a WetLabs ECO-BB3 which was converted to backscattering (b_b_) measurements. Both instruments were manually switched between running 0.2 um filtered sea water and total (“normal”) sea water the rest of the time. Filtered seawater was run before and after each transect and absorption, attenuation, and backscattering during the filtered period were linearly interpolated and subtracted from the total sea water values to get the particulate a (a_p_), c (c_p_), and b_b_ (b_bp_). This setup allows retrieval of particulate optical properties independently from instrument drift and biofouling (Slade et al., 2010). This data was collected using Inlinino (Haëntjens & Boss, 2020), an open-source logging and visualization program, processed using InlineAnalysis (https://github.com/OceanOptics/InLineAnalysis) following (E. Boss et al., 2019). Colored dissolved organic matter (CDOM) was also measured with a Seapoint Ultraviolet Fluorometer logged on a DataQ DI-2108 to Inlinino. (See supplemental material for more detailed processing overview)

These inherent optical properties (i.e., absorption, attenuation, and backscattering) and products calculated from them were used as proxies for a range of particulate properties. γ, which is estimated as the spectral slope of c_p_, is a strong proxy for particle size distribution with a higher γ indicating smaller average particle sizes (Emmanuel Boss et al., 2001). Scattering is driven primarily by particle concentrations, and backscattering is sensitive to both particle refractive index and concentrations, thus the backscattering ratio (i.e. backscattering to scattering) normalizes concentration and varies primarily with refractive index, serving as a proxy for particle composition (Twardowski et al., 2001). A proxy for phytoplankton size (HH_G50) based on anomalous dispersion in the narrow chl-*a* absorption band around 676 nm is also reported (Houskeeper et al., 2020). The last optical proxy we used is phytoplankton pigment concentrations derived from a gaussian decomposition of the particulate absorption spectrum following (Chase et al., 2013). This approach uses a series of gaussians placed at the same location as various pigment absorption peaks and minimizes the difference between a spectrum constructed of these gaussians with the measured spectrum. This gives the approximate concentration of a range of pigments that can be used as a proxy for various phytoplankton taxonomic groups.

### 2.8 Underway Profiling

Profiles were conducted with a Rockland Scientific VMP-250 which collects S, T, chlorophyll-a fluorescence, turbidity (via optical backscatter at 880nm), and turbulence. This instrument was operated in a tow-yo mode on a winch which allowed it to freefall and then be quickly reeled back in and repeated. Profiles were deployed to roughly 100m depth approximately every 500-800m along track. Only downcasts were used.

### 2.9 O_2_/Ar-based Net Community Production

Net Community Production (NCP), a measure of the net production minus net respiration in the system, was estimated by continuously measuring O_2_/Ar ratios in the seawater using Equilibrator Inlet Mass Spectrometry. Briefly, the biological oxygen supersaturation in the mixed layer was calculated following:

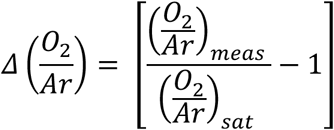

where (O_2_/Ar)*meas* is the ratio of O_2_ and Ar measured in the seawater and (O_2_/Ar)*sat* is the ratio of O_2_ and Ar in the air. Mixed layer depth (MLD) was calculated using a change of 0.1kg m^−3^ from the surface density based on VMP-250 profiles. MLD was interpolated linearly to get to 2-minute average ML. MLD was fairly consistent in the shelf, slope, eddy, and Gulf Stream water types and thus night time MLD was based on day time MLDs and water type. NCP was calculated following:

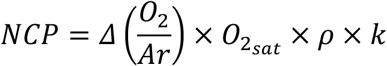

where O_2sat_ represents the saturated oxygen concentration (μmol kg^−1^) (<u>Garcia and Gordon 1992</u>), *ρ*represents the density of the seawater (kg m^−3^), and *k* represents the weighted gas transfer velocity (m d^−1^) calculated following (Wanninkhof, 2014) and (Reuer et al., 2007), with modifications from (Teeter et al., 2018). NCP is reported here in units of mmol O_2_ m^−2^ d^−1^. 6-hour wind speed data were downloaded from the ERA 5 REanalysis dataset (https://www.ecmwf.int/en/forecasts/datasets/reanalysis-datasets/era5). More detailed NCP calculation methods can be found in (Cassar et al., 2009).

### 2.10 ADCP

Ocean velocity data was collected using a Nortek Signature 500 VM Acoustic Doppler Current Profiler. This instrument operates at 500kHz and collects horizontal current vectors and acoustic backscatter. From this, current speed, direction, and shear were calculated along with backscatter strength. Vertical profiles of these measurements were binned in 1m increments from ~1 to 60m depth.

### 2.11 Satellite data

We used multiple ocean-observing satellites to provide context for the vessel-based sampling. Chl-a products were derived from the European Commission Copernicus programmes Sentinel-3’s Ocean and Land Color Instrument (OLCI) and the Ocean Colour Climate Change Initiative’s multi-sensor global satellite chl-a product (Sathyendranath et al., 2019). The OLCI data was used to track chl-a within the eddy over its lifetime. Sea surface temperature products were from the Geostationary Operational Environmental Satellite 16 (GOES-16). OLCI Level 2 and obtained from https://coda.eumetsat.int, OC-CCI chl-a was version 5.0 eight day composites from https://www.oceancolour.org/, and GOES-16 hourly SST was acquired from https://cwcgom.aoml.noaa.gov/erddap/griddap/goes16SSThourly.html.

### 2.12 Delineating the water masses based on Salinity and Temperature

Analysis is primarily based on a continuous view of water properties, given the amount of turbulent mixing occurring along the front, but we also divide water masses into discrete categories of shelf, slope, eddy, and Gulf Stream based on temperature and salinity. That delineation is done by categorizing all water where Shelf < 34 PSU, 35.75 PSU > Slope > 34 PSU, 28.2 C > Eddy > 35.75 PSU, and Gulf Stream > 28.2 °C as shown by the T-S diagram in Figure 4 and 5.

**Figure 5.**
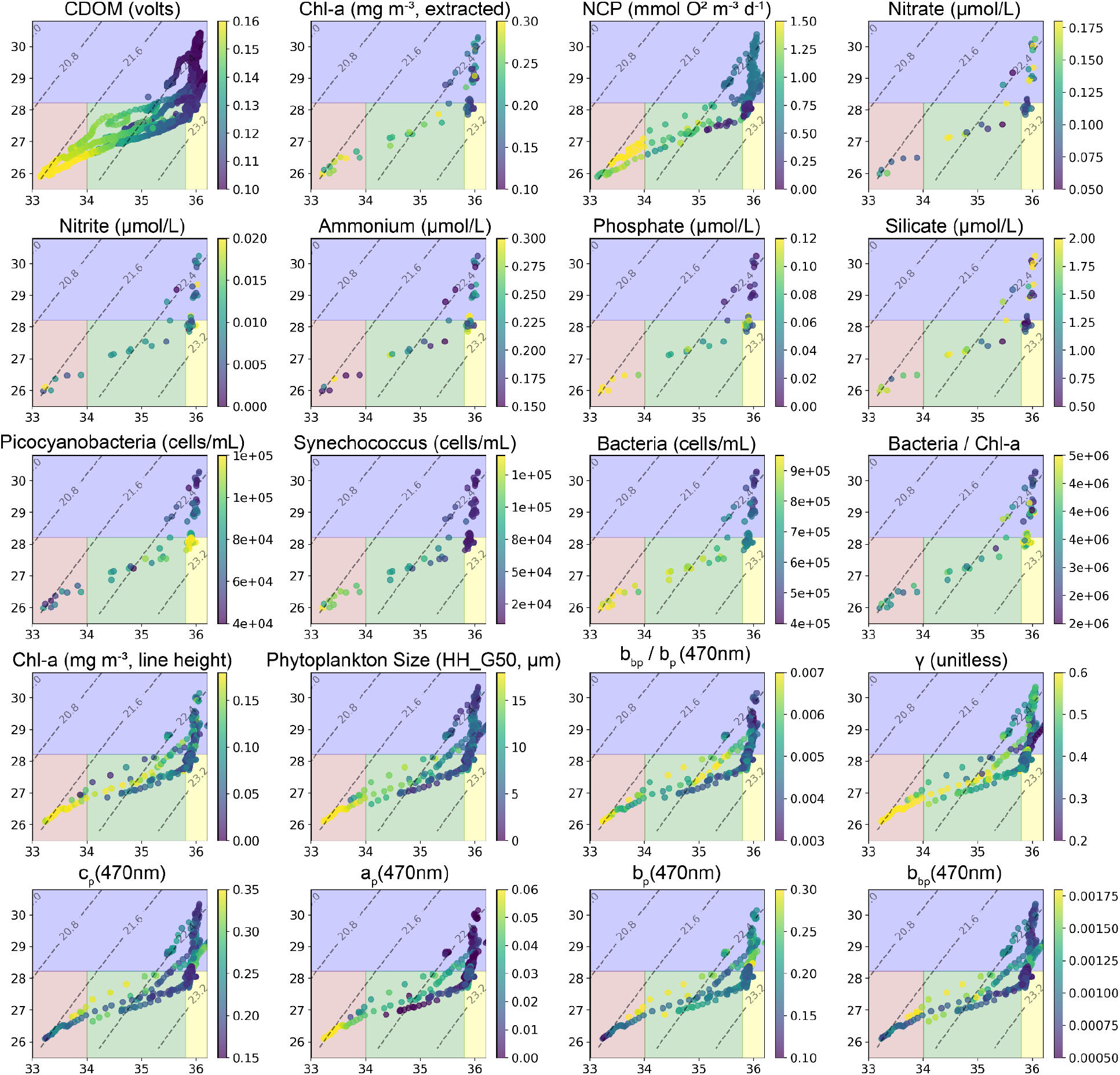
Measured parameters are shown on these T-S diagrams with salinity on the x-axis and temperature on the y-axis. CDOM shows the expected general trend of decreasing dissolved organic matter as you move from the shelf (colder and fresher) to the Gulf Stream (hotter and saltier). While many properties show a general increase or decrease from the Gulf Stream to the shelf - with the eddy simply near the middle or resembling the Gulf Stream properties - a few properties stand out in the eddy from either side. These are namely elevated picocyanobacteria, elevated bacteria/chla ratio, depleted silicate, and depleted nitrate.

### 2.13 Eddy Microbiome

A companion study was conducted which focused on the microbiome of the eddy and adjacent water masses (Gronniger et al., 2023). This companion work ran 16SrRNA gene sequencing on the same water samples as our flow cytometry and nutrient analysis and compares the genetic makeup of microbes across the physical structures we studied. The results of Gronniger et al (2023)’s work are summarized and included in the discussion of this study for context.

## 3 Results

Our data indicate multiple distinct water masses and clear transitions between the shelf, slope, eddy, and Gulf Stream water (Figure S2). While most biogeochemical properties shift gradually and monotonically across water types, some measured parameters cluster into four distinct groups matching the independent discrete T-S based definitions (Figure 5). Salinity and temperature increase from shelf to slope to eddy to Gulf Stream, while CDOM, phytoplankton size, *Synechococcus*, and bacteria all decrease along this continuum. Physically, the eddy water is similar to Gulf Stream salinity, but is slightly cooler in temperature and thus denser. After classifying water masses based on T and S more nuanced patterns are visible (Figure 6). At first glance, the eddy appears to be on the continuum between slope and Gulf Stream water or, at times, indistinguishable from the Gulf Stream. This is true for most of the optical proxies. Measurements of c_p_ are similar in the shelf, slope, eddy, and Gulf Stream waters (Figure S4). c_p_ is dominated by b_p_ in all water masses, though a_p_ is a substantially larger proportion of c_p_ in the 400-500nm range in the shelf compared to eddy and Gulf Stream water (Figure S5) as we might expect given the higher chl-a concentrations on the shelf. The backscattering ratio at 440nm (representative of refractive index and thus composition) generally decreases marginally as we move offshore, indicating slightly more organic relative to inorganic particles. Phytoplankton size (via HH_G50) decreases monotonically with CDOM as we move offshore, though surprisingly γ (negatively correlated with the particle size distirbution) decreases as we move offshore, indicating larger particles offshore compared to inshore, in contrast to HH_G50 trends.

**Figure 6.**
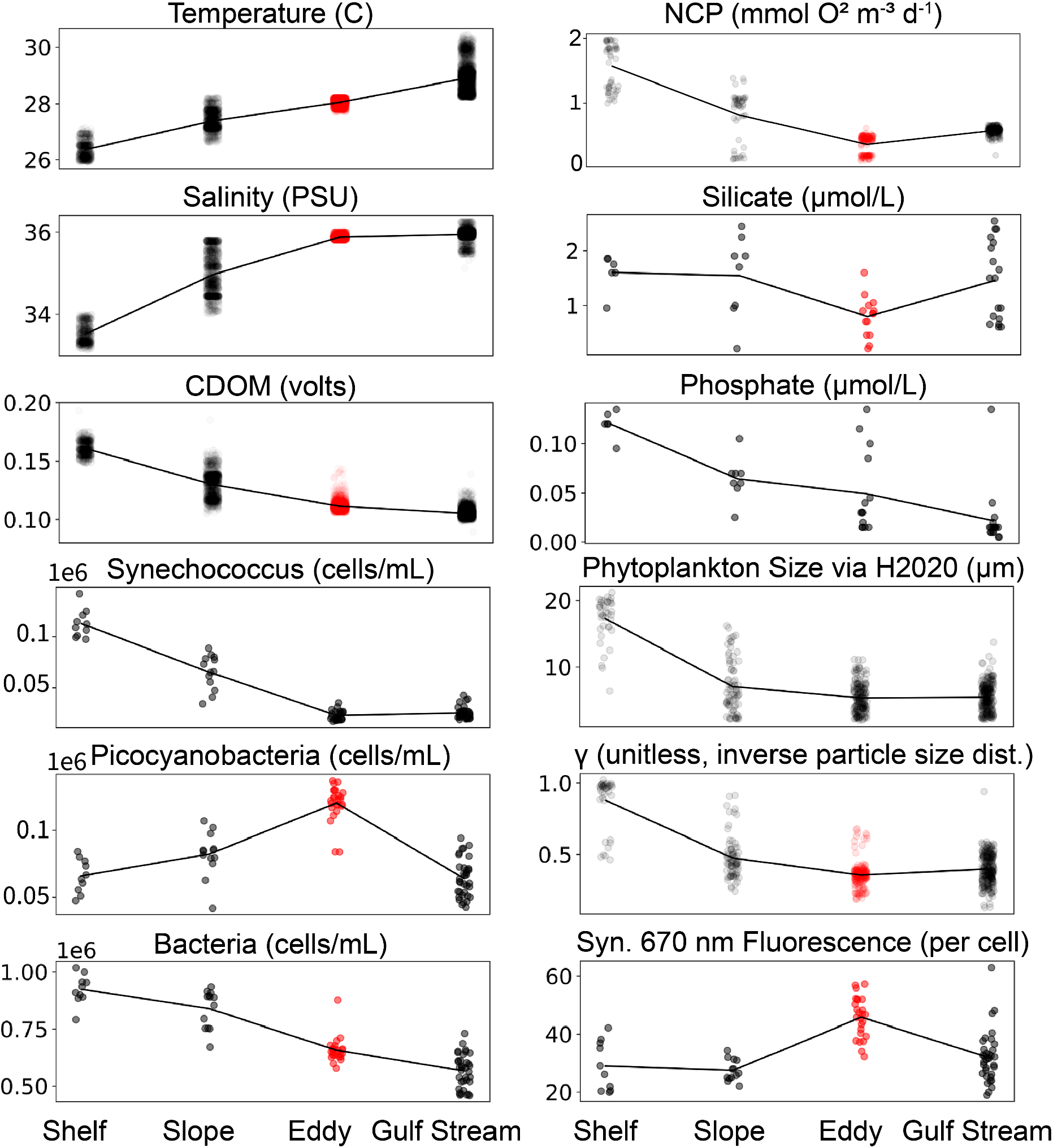
Subset of measured surface variables after classifying water masses into discrete groups (shelf, slope, eddy, and Gulf Stream) based on temperature and salinity properties. Red color in the eddy samples indicates it is statistically different from all other water masses (Welch’s T-test with a p value < 0.05). Some variables are simply a continuum through these water types, monotonically decreasing as T or S increases (e.g. CDOM, bacteria, PO_4_) and some properties are notably different in the eddy and could not be a simple product of mixing (e.g. picocyanobacteria, NCP, fluorescence per cell). A few variables including NO_2_, NH_4_, and NO_3_ are not statistically different between any water mass (e.g. NO_3_ ~0.1umol/L in all water masses). Not shown here, but fluorescence per cell is nearly identical for non-phycoerythrin containing picocyanobacteria.

A few properties however reveal the eddy as distinct from the Gulf Stream, with depleted nitrate and silicate compared to the Gulf Stream water, enhanced non-phycoerythrin containing picocyanobacteria compared to all other water masses, the highest bacteria to chl-a ratio, and lowest volumetric NCP (Figures 5 and 6). In the eddy c_p_(460), and b_bp_(440), and γ are marginally lower compared to Gulf Stream (Figures 5 and 6).

Profiling data shows a MLD of ~10m for shelf and slope water with eddy or Gulf Stream water underlying that shallow shelf/slope water and a second thermocline around ~30-50m. The MLD for the eddy ranges from approximately 30-50m and Gulf Stream is the same range from 35-55m. Calculating MLD at the front at these scales is often problematic given the intense mixing and turbulence across the front, so these exact depths should be interpreted with caution. We observe a fairly consistent cold and fresh intrusion (Figure S8) moving from the shelf to the Gulf Stream and converging with the Gulf Stream that could be of Arctic or northern slope origin given the temperature (<15C) and salinity (33-33.5 PSU, comparable to shelf). While not the focus of this study, this cold and relatively fresh layer has intense structuring of both chl-a fluorescence and echosounder volume backscatter.

ADCP data supports conclusions from profiling and surface categorizations, showing faster current speeds in the Gulf Stream water vs eddy water and cyclonic rotation in Transect 9 (T9, Figure 8). Echosounder data from the ADCP shows fairly consistent backscatter responses in Gulf Stream and Eddy water and much stronger backscatter in the shallow shelf water (Figure 9). This acoustic backscatter data also shows a thin layer at the thermocline of the eddy that isn’t apparent in the Gulf Stream water (Figure 8).

Looking back over the satellite record indicates a monotonic decrease in chl-a at each observation time step from eddy formation until our survey work (Figure 7).

**Figure 7.**
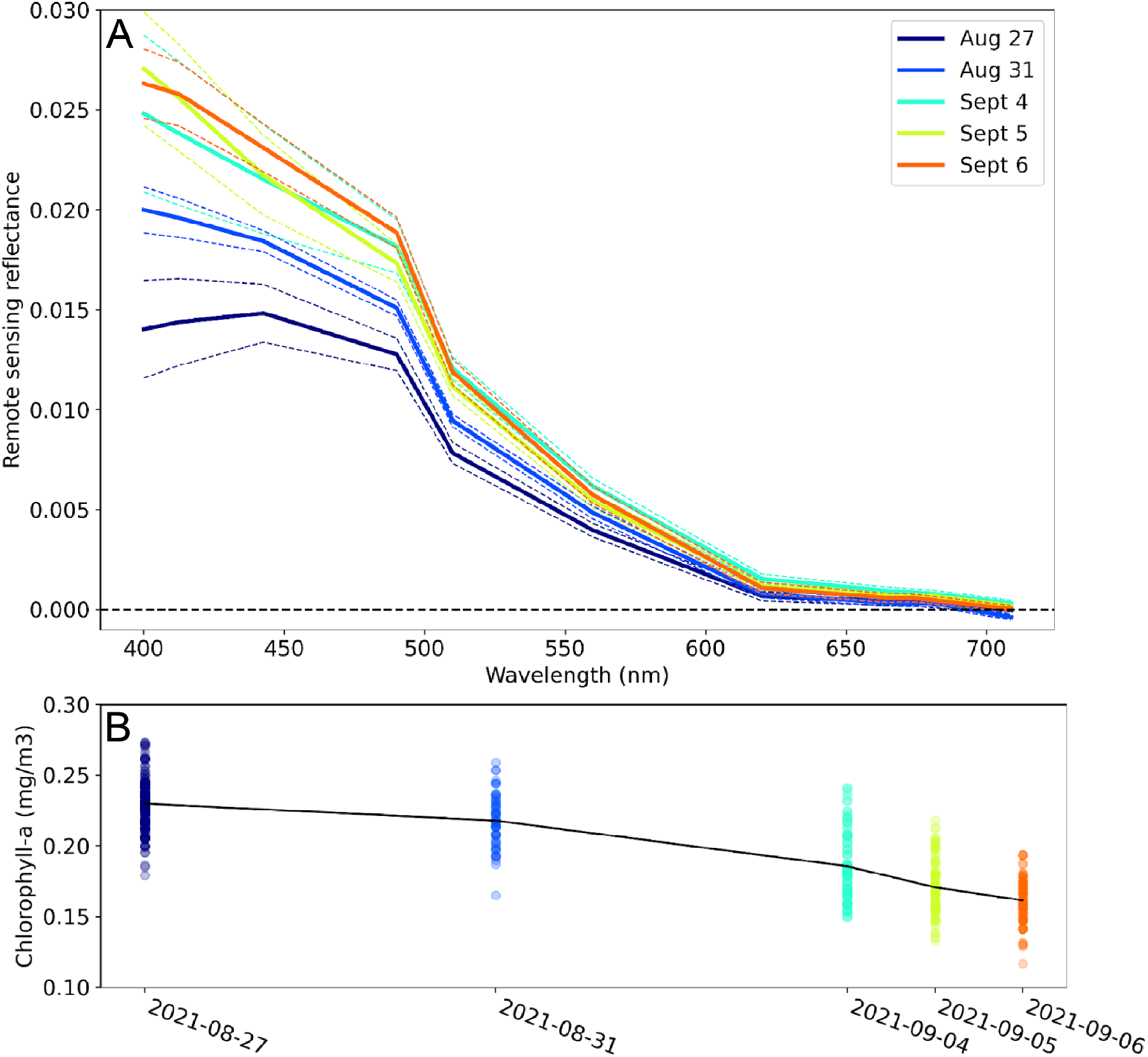
A. Shows median Rrs spectra of the eddy from the Sentinel-3 OLCI sensor over the lifetime of the feature. This shows an increase in the blue wavelengths over time from the formation of the eddy up until the last day it was coherent and not cloudy. Dashed lines show standard deviation of spectra within the eddy. Black dashed line demarcates and Rrs value of 0. B shows chl-a concentration (calculated via the Hu et al 2012 ocean color index) decreasing monotonically throughout the observational period, the colors represent the same dates as in the top panel and the black line connects the means of each time step. All satellite data used in this figure shown in Figure S1.

The VMP data reveals the physical structure across all transects, with a general trend of fresher and less dense shelf/slope water sitting on top of the eddy/Gulf Stream water in a thin layer 5-20m and under this water is eddy or Gulf Stream water (Figure S8). Then where the shelf/slope water ends at the surface the eddy/Gulf Stream water reaches the surface. The peak in chl-a fluorescence from VMP data is typically around 30-50m, just below the thermocline.

On the last transect of the cruise, T9, the ADCP indicates we crossed the entirety of the eddy near the middle of the feature and we see strong evidence for continued rotation (Figures 8 and S7), though at relatively low speed (10-40 cms^−1^) compared to the Gulf Stream transects (T2, 150 cm s^−1^). The ADCP also shows clear structuring of acoustic backscatter (representative of zooplankton and small fish). The highest values of acoustic backscatter are in the shelf water. Within the Gulf Stream and eddy water we observe high backscatter from the surface down to

**Figure 8.**
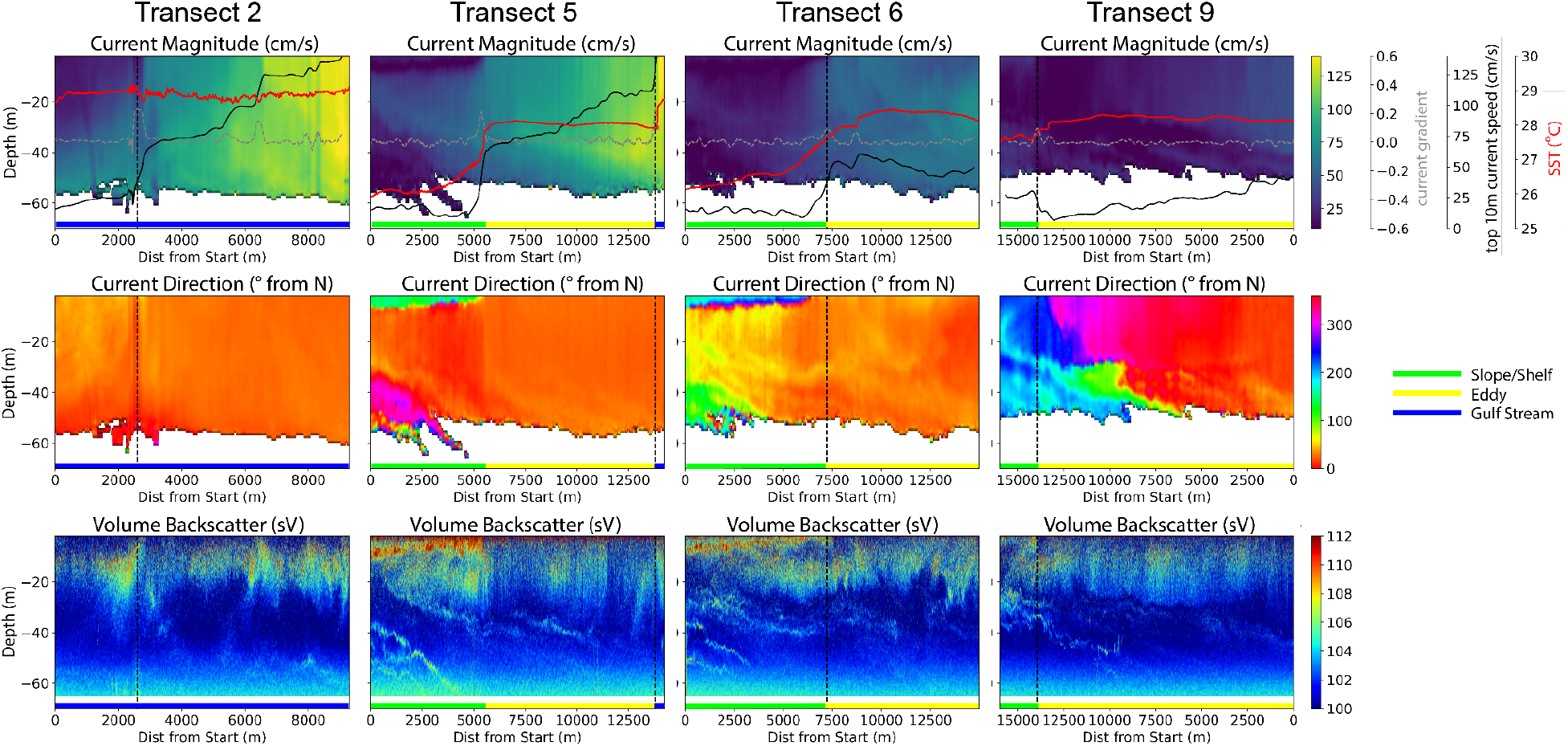
Current magnitude and direction from the ship-based ADCP are shown in the top two rows and the third row shows volume backscatter from the echosounder on the ADCP. The vertical dashed black line is the front as defined by the peak gradient in current, the red line is sea surface temperature as measured by the ADCP, the black line is current speed, and the grey line is the current gradient. Current magnitude (top panel) shows clear delineations between shelf water and the eddy and the eddy and the Gulf Stream. The acoustic backscatter data (bottom panel) shows the most intense backscatter in the shelf water with some possible structuring from the eddy and strong thin layers from the cold water intrusion below the eddy. Note T2 data is only from half of the transect due to an instrument error.

~35 m (effectively the mixed layer) and then comparatively low backscatter from ~35 to ~60 m. From 60m and deeper the data is overwhelmed by noise. As these different water masses are layered physically, so are the backscatter responses. For example in Transect 6 (T6), around 4000 m into the transect laterally, there is highly scattering shelf water from 0-8 m, then eddy water with backscatter that eventually decreases further around 30 m, followed by 3-4 thin backscattering layers of 1-3 m, first at the thermocline where the eddy ends and then above and below the cold water intrusion.

## 4 Discussion

Given the frequency of frontal eddies traveling along the Gulf Stream past Cape Hatteras, it was our goal to understand their state after passing Cape Hatteras and the possible biogeochemical impacts in the MAB. Using this information, we assess if they could help explain the productivity of “the Point” of Gulf Stream separation from the continental shelf and broadly how they may be impacting the MAB.

### 4.1 Eddy history

The satellite record (Figures 7 and S1) indicates a formation and evolution of the observed frontal eddy in line with observations from previous studies and recent modeling (Glenn & Ebbesmeyer, 1994b; Gula et al., 2016; T. N. Lee et al., 1991) and suggests that this eddy is a typical example of a frontal eddy in this region. The eddy formed off the Charleston Bump (Figure 1) driving a chl-a and SST anomaly within a Gulf Stream meander. The satellite data does not show a time-evolving surface bloom compared to the initial shelf waters, but the eddy does initially have higher surface chl-a than the Gulf Stream water. Over the course of two weeks there is a gradual decrease in chl-a either from die off or mixing with Gulf Stream water. This lack of a satellite-visible increase in chl-a when compared to the initial shelf conditions could be due to enhancement of growth occurring at a depth below that detectable by satellite, from a concomitant increase of grazers, or limited upwelling. We suggest it is a combination of enhancement at depth and then grazers based on previous work on frontal eddies in the SAB and across the globe (Glenn & Ebbesmeyer, 1994a; Gula et al., 2016; Kasai et al., 2002; Yoder et al., 1981). The satellite record shows the eddy had been coherent for two weeks at the time of our in-situ observations.

### 4.2 Physical status during cruise

During our cruise the eddy was still mostly coherent with consistent physical properties across the feature and continued cyclonic rotation, which, while weak, would likely be driving a negative SSH anomaly and upwelling within the eddy, possibly still enhancing productivity. However, higher resolution and deeper sampling within the water column would be needed to reveal the eddy’s core and any stimulation of primary producer biomass.

On T6 we observe what is likely the remains of the frontal eddy’s warm streamer that has been pulled into the body of the eddy (Figure S2 for a detailed description of each transect along with S and T). It is apparent from the surface to ~30m depth where temperature and salinity are higher than the rest of the eddy and cell counts from flow cytometry are closer to the Gulf Stream. Salinity and temperature in this warm streamer are intermediate between Gulf Stream and eddy, likely representative of the intense mixing that occurs in frontal eddies (Gula et al., 2016), especially past Cape Hatteras where they begin to dissipate.

### 4.3 Ecological observations

In spite of predicted upwelling at depth, in general the eddy has more oligotrophic conditions compared to the other water masses, with lower silicate and and NO_3_ in the eddy compared to Gulf Stream and slope waters and enhanced non-phycoerythrin containing picocyanobacteria (a group which contains *Prochlorococcus*) compared to other water masses (Figure 6). While cyclonic eddy dissipation will relax the isopycnal uplift, causing downwelling (see Figure 5 in McGillicuddy, 2016), it is unclear exactly how this process works within frontal eddies and if the timeline matches for this to help explain the more oligotrophic conditions. Combined with the nutrient data which reveals lower silicate and NO_3_ in the eddy compared to Gulf Stream and slope waters, this suggests a shift to a microbial loop dominated system. The highest bacteria to chl-a ratio occurs within the eddy, suggestive of post-bloom conditions (Buchan et al., 2014). NCP is another variable in support of post-bloom conditions with the lowest volumetric NCP being measured in the eddy, slightly lower than the Gulf Stream conditions and substantially lower nearer the core of the eddy on the final study day (Figure 5).

A companion study (Gronniger et al., 2023), focused on the eddy microbiome, was undertaken on the same flow-through surface samples and similarly reveals differences in microbial community composition between water masses. This companion study found that the eddy harbored higher abundance of *Prochlorococcus* and lower abundances of *Synechococcus* and *Pelagibacteraceae*. However, despite differences between the Gulf Stream and eddy microbiomes, including higher abundances of *Prochlorococcus* in the eddy and higher abundances of the nitrogen-fixer *Trichodesmium* in the Gulf Stream (see also Figure 6), the eddy is most similar to the Gulf Stream relative to the other water masses, suggesting that assembly of the eddy microbiome is primarily determined by environmental filtering in these warm, low nutrient waters. Gronniger et al (2023) clustered the microbiome data and compared this to the physical delineations which showed that the discrete physical classes used here do not fully reflect microbiome clusters, likely due to the intense submesoscale dynamics in this region.

Fluorescence per cell via flow cytometry is highest in the eddy and in frontal regions (Figure 6). This was not our expectation considering higher fluorescence per call can be indicative of either more nutrients or more pigments per cell. We thus would expect the slightly lower nutrient conditions in the eddy to result in lower fluorescence per cell. Because higher fluorescence per cell can also indicate cells physiologically acclimatized to lower light levels, this may indicate mixing from depth in these regions or higher nutrient uptake rates, not reflected in free nutrient concentrations (Figure 6).

Acoustic backscatter shows substantial structure in all water masses (Figure 8). As expected, the shelf water, which had the highest chl-a values, has the highest acoustic backscatter. The Gulf Stream and eddy water are not statistically different with respect to backscatter (Figure 9), though in all eddy transects there is a clear thin layer of enhancement in backscatter around the thermocline of the eddy that does not appear in the Gulf Stream water (Figure 8). Given the limited number of samples, it is possible this difference is related to other factors and not the dynamics of the eddy itself. The cold water intrusion also appears to drive thin layers in acoustic backscatter above and below the feature. Interestingly, the peak in chl-a fluorescence from the VMP data is sometimes shifted down by ~15m from the acoustic backscatter peak. This could be due to grazing decreasing biomass and thus fluorescence, non-photochemical quenching of fluorescence in the better-lit upper layer, the relatively high 1000kHz sampling frequency not picking up on the grazers consuming the phytoplankton, or actual partitioning in habitat. While some combination of these various factors is the most likely explanation, this is nonetheless a surprising finding. It is unlikely quenching is drastically different at 35m vs 45m and could suggest that in this region standing phytoplankton biomass (here proxied by chl-a) is not a good representation of productivity.

**Figure 9.**
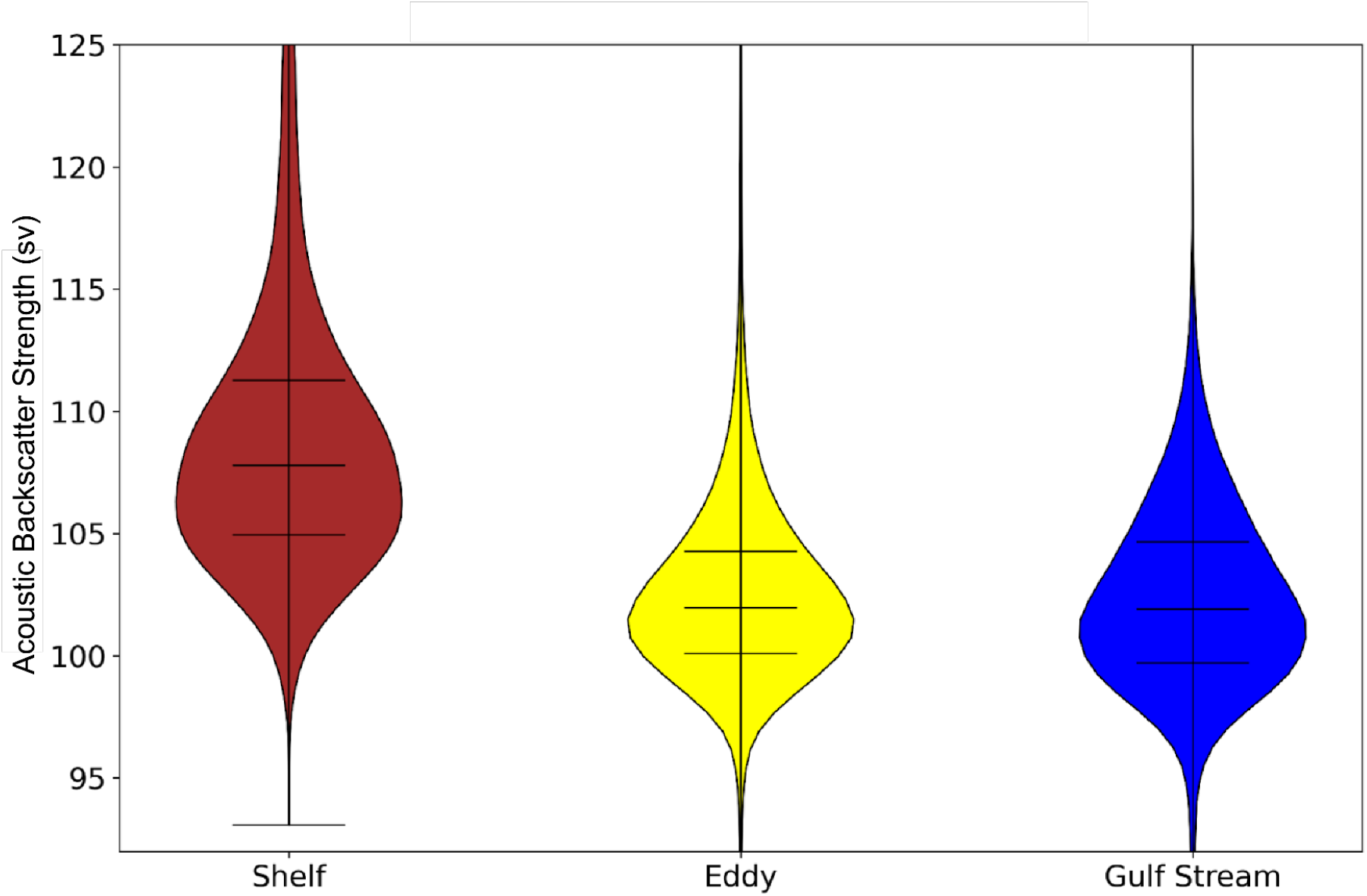
Acoustic backscatter strength across water masses from the ADCP. This is the distribution of acoustic backscatter in the mixed layer for the shelf (down to 10m), eddy, and Gulf Stream water (both averaged down to 40m). Means are respectively 108.97, 102.56, and 102.34 for the Shelf, eddy, and Gulf Stream water. ADCP measurements were not collected in the slope water and thus are not represented here.

### 4.4 Particle Size Surprises

One interesting note from this work is the divergence of γ (proxy for mean particle size) from typical patterns. γ, which is sensitive to particles in the size range from approximately 3-30 μm (Emmanuel Boss et al., 2001), generally corresponds to chl-a concentration, based on the commonly found relationship that higher chl-a corresponds to larger phytoplankton. Thus, we would expect a decrease in mean particle size as we move from coastal water (high chl-a and larger particles) to oligotrophic water (low chl-a and smaller particles) (Emmanuel Boss et al., 2013; Buonassissi & Dierssen, 2010). In this study, we instead see an increase in particle size and a decrease in chl-a as we move from shelf water further offshore into the eddy and Gulf Stream (Figures 5 and 6).

In general, most particles in the open ocean are phytoplankton or detritus. The observed increase in the mean particle size offshore is unlikely to be indicative of phytoplankton based on HH_G50 which shows decreases in phytoplankton size offshore - following expectations of a positive relationship between chl-a and phytoplankton size. The particles driving γ are therefore unlikely to be live phytoplankton. The backscattering ratio decreases marginally as we move offshore suggesting the particles influencing γ are relatively more organic, just not chl-a containing. Another relevant piece of evidence, b_p_ is not only spectrally flatter in the Gulf Stream (driving c_p_ to be flatter and lower γ), but actually higher overall, indicating higher particle concentrations.

So more particles, larger average particle sizes, and particles that are similarly or relatively more organic than those inshore. This leads us to the conclusion that it is either due to an enhancement of small heterotrophs or possibly detritus and non-algal particles within the eddy and Gulf Stream waters. This could be suggestive of grazer enhancement in the eddy, though again this is not specific to the eddy. It could also be due to an accumulation of buoyant particles near the front.

The relative consistency of γ and HH_G50 between eddy and Gulf Stream waters suggests whatever process is influencing this divergence from expectations is not an eddy specific phenomenon, or if it is, that the resultant particulate impact is mixed into Gulf Stream waters and is possibly persistent between eddies.

### 4.5 Impact of frontal eddies on grazers

While relevant studies investigating consumer response to frontal eddies are rare, previous work shows that copepods and doliolids increase substantially in response to phytoplankton enhancement in frontal eddies. In this case these zooplankton exhibited minimal vertical migration, with maxima near the thermocline, suggesting they would likely be entrained in the frontal eddy and move with it north of Cape Hatteras (Paffenhöfer et al., 1987). The timeline of this increase in zooplankton is variable across taxa, but in this work, the authors found zooplankton increased dramatically from five days to two weeks after peak chl-a concentration, and peak chl-a concentration was approximately a week after eddy formation. Similar work has shown that copepods, which are slower to respond, may persist for a few weeks, and doliolids which respond within days, were also found to decrease faster, persisting for 7-9 days (Deibel, 1985). This work was done on the southern half of the SAB (with eddy generation around Florida and dissipation just upstream of the Charleston Gyre) and may have different characteristics than the northern section of our study, but if we assume similar responses, zooplankton abundance should peak right when the eddy is passing Cape Hatteras.

Our acoustic data does not support an enhancement of zooplankton and small fish biomass in the eddy compared to Gulf Stream water at the time of our measurements (Figure 9). The variability of some transects adds substantial uncertainty to these comparisons and we don’t have a “pure” Gulf Stream endmember for comparison. Modeling shows substantial diapycnal mixing between these eddies and the Gulf Stream (Gula et al., 2016), possibly contributing to the lack of clear distinction in acoustic backscattering between these water masses. We do see structuring of acoustic backscatter (i.e. zooplankton and fish) with a clear partitioning at the thermocline of the eddy and an increase just below it which is not observed in the Gulf Stream data. This could be an indicator of upwelling and higher growth. Adding to the complexity we observe a large vertical migration via acoustic backscatter into the shelf and eddy water from around 250 m depth. Considering our eddy has spent two nights off the shelf there is a possibility a large amount of the potential grazer biomass enhancement has already been consumed by fish in this migrating layer. To fully parse this out would require a Lagrangian observation approach over the full lifetime of the eddy.

In much of the previous work on frontal eddies the assumption was that these eddies decayed onto the outer shelf of the SAB, and an ongoing question was that if there is such a substantial input of nutrients due to upwelling why is the outer shelf of the northern SAB not more productive in higher trophic levels? We suggest based on our work and previous studies that a large amount of this new production could be moving into grazer biomass, and while some of this may occur on the outer shelf edge in the SAB, these features are still typically being carried by the Gulf Stream into the MAB where they could contribute to the high productivity of “the Point”. Even where there are subsurface intrusions far up onto the shelf of the SAB due to frontal eddy upwelling this water is quickly entrained into the Gulf Stream’s northward flow, and while some of this may stay on the SAB shelf long enough for the grazers to die off or be consumed themselves, most of it is north of Cape Hatteras within 1-2 weeks, shown both by our work and previous studies (Glenn & Ebbesmeyer, 1994a). This is a match in time for grazers to have bloomed following the initial phytoplankton bloom and occurs frequently enough to sustain elevated levels of secondary consumers. Previous work that did consider the entrainment of carbon into the Gulf Stream from these processes framed it as a generic export process rather than explicitly a potential enhancement of zooplankton being delivered directly into the mouths of hungry fish in the MAB e.g. (T. N. Lee et al., 1991). In this sense the northern SAB could be thought of as the breadbasket of the southern MAB with frontal eddies providing the fertilizer.

### 4.6 A refined conceptual model

Using insights from this work we both place our study into the context of the initial conceptual model (Figure 3) and refine that model based on our new understanding.

We begin with the meander at the Charleston Bump which drives the Gulf Stream further offshore. Due to this injection of vorticity the Charleston Gyre exists, and periodically (~3-7 days) drives the creation of a frontal eddy and a warm streamer of Gulf Stream water which encloses this newly formed frontal eddy, partitioning it from the shelf water. We have no in-situ data during this period, but previous work all supports upwelling due to the cyclonic rotation and isopycnal uplift as well enhancement (compared to Gulf Stream water) from trapping during eddy formation (Glenn & Ebbesmeyer, 1994b; T. N. Lee et al., 1991; Yoder et al., 1981). This combination of productive shelf water and upwelling creates a chl-a anomaly well above the mean. Upwelling slowly decreases as the eddy dissipates energy as it moves downstream (Gula et al., 2015). The enhancement in phytoplankton growth enables an increase in grazer populations, which peaks between Cape Lookout and Cape Hatteras, and sustains an enhanced level of secondary consumers, possibly contributing to the high level of top predator populations in this region. By the time the eddy is just north of Cape Hatteras the physics may still be driving some upwelling but the phytoplankton community has switched to gleaners, possibly in a microbial loop. This is where we place our study, around the secondary consumer peak and after phytoplankton biomass has shifted below the baseline. After this shift to smaller cells such as *Prochlorococcus*, the transfer efficiency to fish and top predators is likely to be substantially lowered, adding on to the decrease in available phytoplankton (Eddy et al., 2021).

Our data shows the eddy from this study has depleted Si compared to Gulf Stream and slope water suggesting a possible recent diatom bloom. It has the highest ratio of bacteria : chl-a and lowest NCP again suggestive of post-bloom conditions. The eddy also has a higher ratio of mean particle size : chl-a indicating either an enhancement of micro grazers along with the picocyanobacterial or more detritus per unit chl-a. Both flow cytometry and molecular sequencing show a different community composition dominated by picocyanobacteria and particularly *Prochlorococcus* - again suggestive of a switch to the microbial loop. Thus we hypothesize the eddy arrives at Cape Hatteras with enhanced grazer biomass due to the previous phytoplankton bloom and that these grazers are consumed by the large migrating layer we observed.

Within the SAB frontal eddies are thought to have a dominant impact on nitrogen fluxes on the outer shelf (T. N. Lee et al., 1991) - with a huge impact on the ecology of the SAB outer shelf. Based on our observations, frontal eddies do not appear to supply phytoplankton into the MAB, but are possibly an efficient shuttle of grazers into the highly productive southern MAB which is thought to have some of the highest marine mammal diversity in the world and is a productive fishing ground (Byrd et al., 2014).

Given the complexity and dynamic nature of this region we can only hypothesize the impact of frontal eddies on the overall ecosystem of the SAB and MAB (see supplemental material for extended caveats). In future studies we need a Lagrangian survey approach over the entire lifetime of the eddy. With acoustic backscatter and discrete net tow samples we could identify the density and type of zooplankton within the eddy. In this work we focused on surface samples but a large component of the frontal eddy driven enhancement may be below the thermocline as it is lifted into a more favorable light environment. A fleet of autonomous assets would facilitate a more synoptic view of the eddy. A survey design similar to (Zhang et al., 2021) with AUVs and gliders to assess the full physical and biogeochemical status could enable a test of our hypothesis. This could be an ideal exercise for high resolution modeling to examine the MAB ecosystem with and without frontal eddies.

### 4.7 Broader Context

A similar process may be happening just upstream (south) of the Charleston Bump, where the timeline matches up for the enhancement of grazers after frontal eddy enhancement from Miami to Charleston and possibly complemented by a longer duration stay in the Gyre. Even more generally, anywhere a western boundary current follows the continental shelf and the system is nutrient limited frontal eddies may be a reliable mechanism for increasing primary production and transfer efficiency to higher trophic levels.

## 5 Conclusions

We show in this work that frontal eddies do in fact often get advected north past Cape Hatteras, can still have cyclonic rotation, and have nutrient profiles and phytoplankton community compositions distinct from adjacent water masses even in dissipation. We hypothesize they may be well timed to supply zooplankton to secondary consumers between Cape Lookout and Cape Hatteras. Synthesizing previous literature we share a simple conceptual model for the ecosystem impact of frontal eddies and place our study within that model and add refinements. Given the frequency of frontal eddies moving along the Gulf Stream this may be a mechanism for substantial enhancement of phytoplankton, zooplankton, and secondary consumers - particularly at the highly productive separation point of the Gulf Stream jet. We suggest further work on frontal eddies in the Gulf Stream and other western boundary currents to see if this is a general phenomenon.

## Acknowledgments

Funding support was provided by National Aeronautics and Space Administration (NASA) Future Investigators in NASA Earth and Space Science and Technology (FINESST) #80NSSC19K1366 (Ocean Biology and Biogeochemistry program), the Zuckerman STEM Leadership Program, the American Society for Photogrammetry and Remote Sensing’s Robert N. Colwell Memorial Fellowship to PCG. R/V Shearwater ship time was supported by Nicholas School of the Environment and Duke University Marine Lab donors through a student research grant to PCG. We appreciate Emmanuel Boss for sharing instruments for the optical flow-through data. Finally the authors thank the crew of the R/V Shearwater, Matt Dawson, Tina Thomas, Zach Swaim, and John Wilson for making much of this data feasible and enjoyable to collect.

## Open Research

The code to recreate this analysis are available at https://doi.org/10.5281/zenodo.7685135 (Gray, 2023) and all data used in this study is available at https://doi.org/10.5281/zenodo.7680135 (Gray et al., 2023). All code is shared with an MIT License for free reuse. We have provided multiple Jupyter Notebooks that go from raw or initially processed data to the nearly complete figures shown in the paper. The python environment used can be easily and exactly reproduced using the pangeo-notebook a Docker image https://github.com/pangeo-data/pangeo-docker-images/tree/master/pangeo-notebook as detailed in the Github repo.

