## Supplemental Material for "The Impact of Gulf Stream Frontal Eddies on Ecology and Biogeochemistry near Cape Hatteras"

**Additional Optical Processing Information**

All attenuation (c_p_), absorption (a_p_), and backscattering (b_bp_) spectra were manually quality-controlled to remove noisy or erroneous data (e.g. bubbles, bad filtered seawater measurements). For each minute of the total sea water measurement, the signal between the 2.5th and 97.5th percentiles were averaged and standard deviation was calculated to quantify uncertainty. Values beyond 2.5th to 97.5th percentiles were removed to filter noisy spikes due to bubbles which are known to cause erroneous ACS measurements. All a_p_ and c_p_ spectra were unsmoothed following the method in Chase et al. (2013) and a residual temperature and scattering correction was applied using the temperature and salinity dependencies from (Sullivan et al. 2006). Following (Sullivan et al. 2013) χ=1.076 was used to convert from scattering at a single angle (nominal angle 124°) to the particulate backscattering coefficient (b_bp_).

**Extended Caveats**

Given that the eddy-induced upwelling is typically around the thermocline at 30-50m depth, the primary impact of this eddy is at depth and most of our measurements are at the surface. While we do expect these properties are being mixed throughout the mixed layer, future work is warranted taking similar measurements as done here but across vertical sections of frontal eddies. In this work we opted for surface measurements in order to capture a more synoptic view given the rapid currents, but measurements at depth are a missing and possibly critical part of this story.

Our survey locations were not evenly spread among the eddy, our assumption is that the waters in the eddy are reasonably consistent but we can’t confidently demonstrate this and in fact satellite data shows that intense mixing is going on with rich submesoscale variability primarily at the Gulf Stream edge of this eddy. Particularly the nutrient profile around the cold core, if it is still upwelling, may be quite different. Given that the eddy is partially sheared apart it is challenging to differentiate exactly where we are in the eddy structure and a portion of our data is on the far downstream edge of it.

Our discrete categories are based on the assumption that we have representative “endmembers” of the shelf, slope, eddy, and Gulf Stream. This could be problematic since the spatial scale of this work is relatively small (~10km transects) and these water masses are all actively mixing and more of a continuum with no “pure” endmembers. This is particularly true of the Gulf Stream and slope waters. In the Gulf Stream water we *a priori* expected the most consistency, but properties are fairly variable, likely due to entrainment of shelf and slope water into the Gulf Stream. The slope water is inherently a transitional water type and thus a mix of shelf and Gulf Stream. And our eddy has been mixing intensely with the Gulf Stream for two weeks and many of our eddy samples are from the downstream edge where we expect intense strain is occurring driving mixing. Additionally the Gulf Stream samples took place further upstream, just as the eddy was moving into the area (as evident in the “shattered front" temperature variability in Transect 2), in shallower water where the bottom might be impacting our measurements.

A concurrent study with samples taken at the same stations as presented here [Gronniger et al 2022] was able to discriminate four distinct clusters via 16S rRNA gene sequencing and while very similar these did not exactly match up with the physical clusters based on salinity and temperature. This was particularly true of the eddy and Gulf Stream water where there has been intense mixing and the warm streamer has possibly been pulled back into the eddy and mixed in. This suggests some of those comparisons need to be taken with a grain of salt, and suggests that possibly the best timing to inspect grazer enhancement is actually just before passing Cape Hatteras or even Cape Lookout when the eddy is more coherent.

One of our main ecological questions, whether frontal eddies enhance grazers for the MAB, is not convincingly answered by our data. More work will need to be conducted in order to have a definitive answer. For the time being it is certainly one of a few options in the set of frontal eddy impacts based on the processes we have observed and could be of substantial import to the MAB.

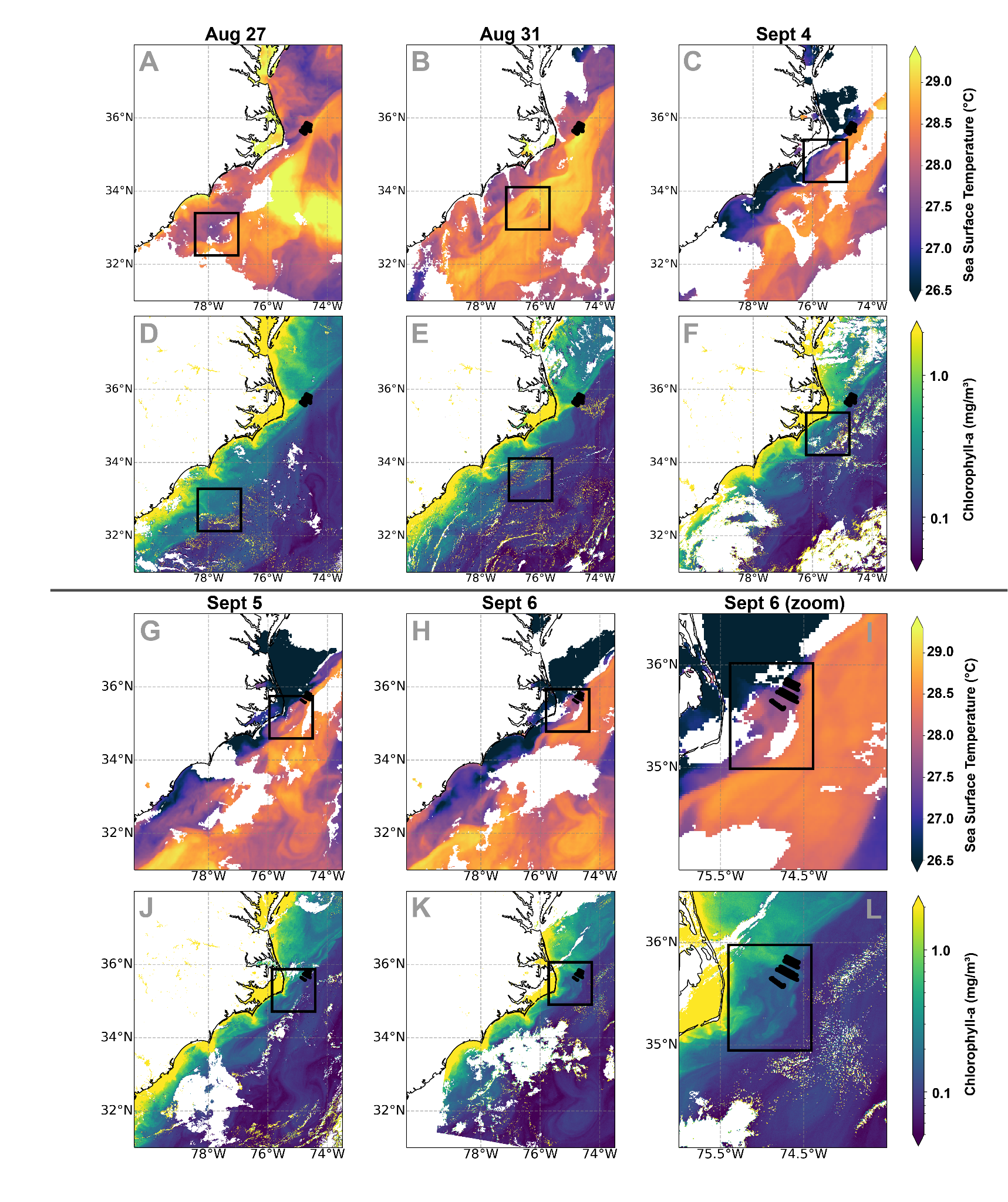

**Figure S1**. Satellite overview of the observed frontal eddy. Black dotted lines are all 9 transects from this study. Panels A-C and G-I show sea surface temperature and panels D-F and J-L show chlorophyll-a. Black boxes outline the eddy location across five days from August 27th to September 6th. SST is from GOES-16 and chl-a is from Sentinel-3’s OLCI sensor.

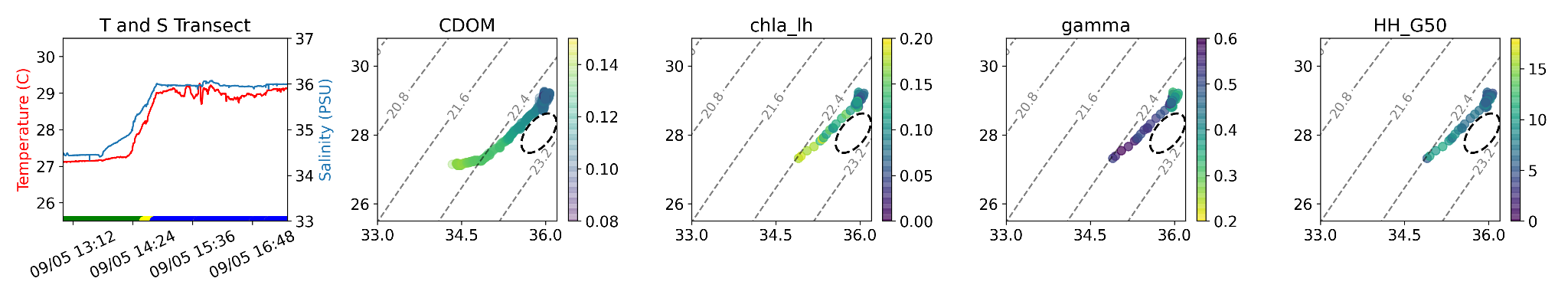

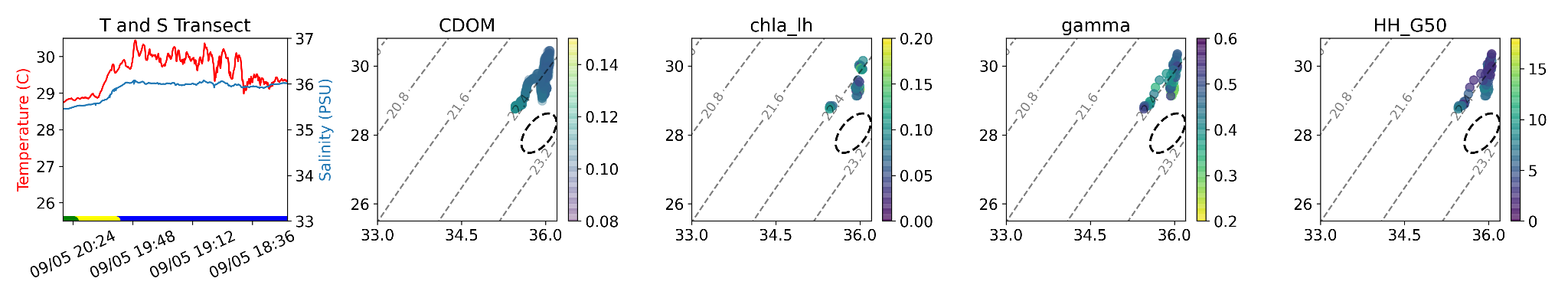

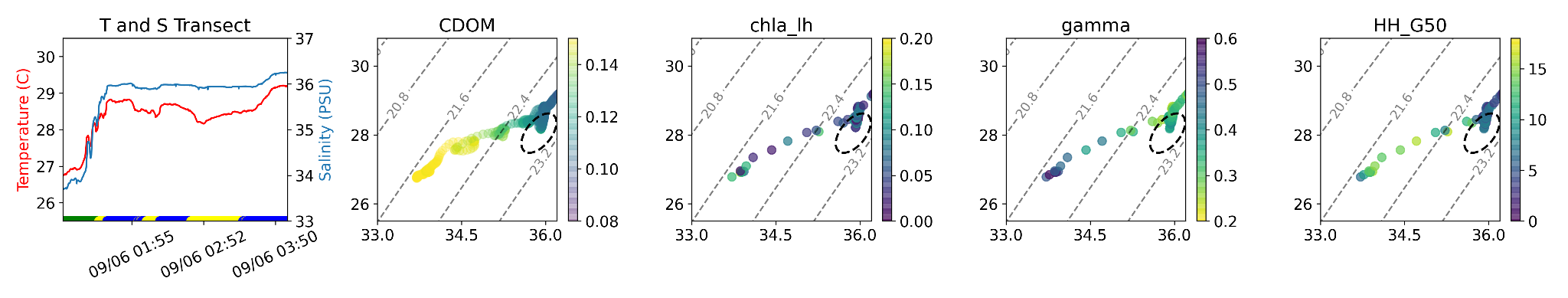

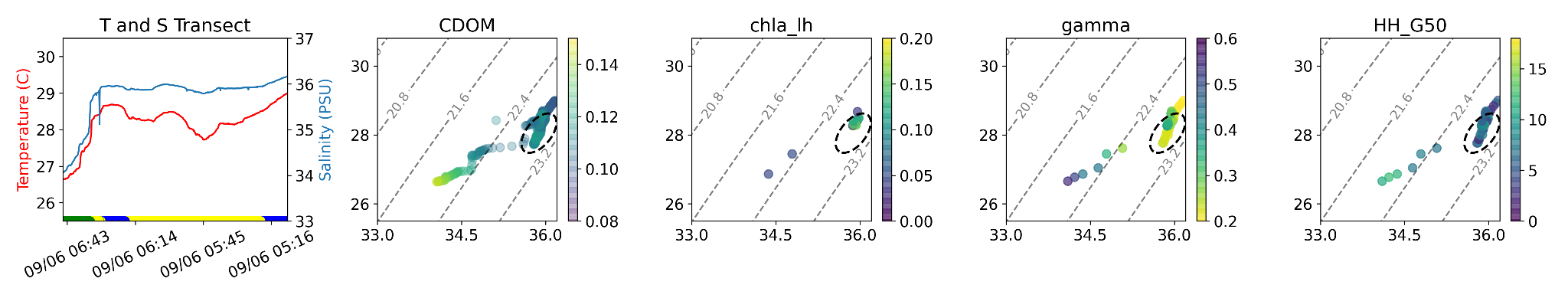

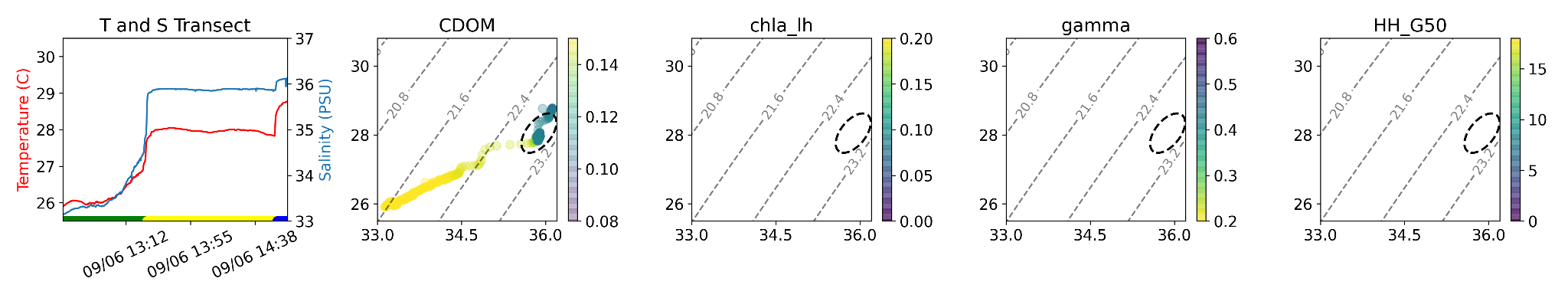

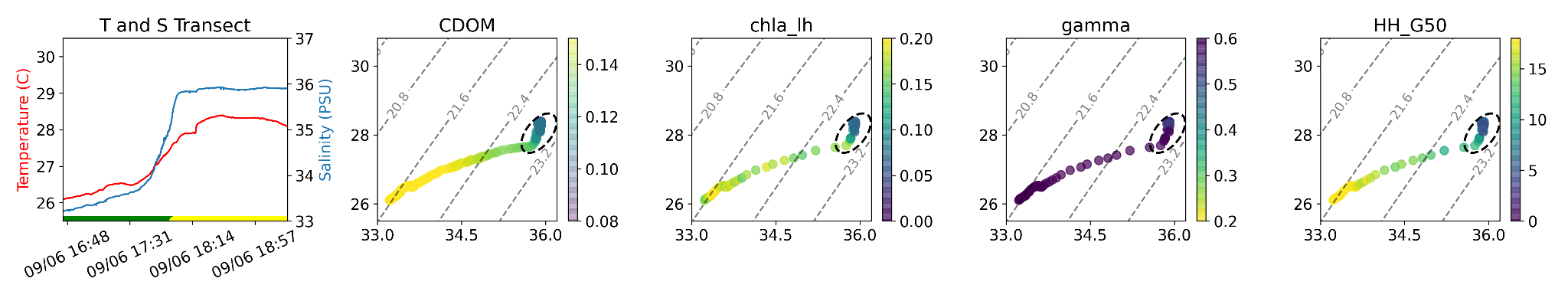

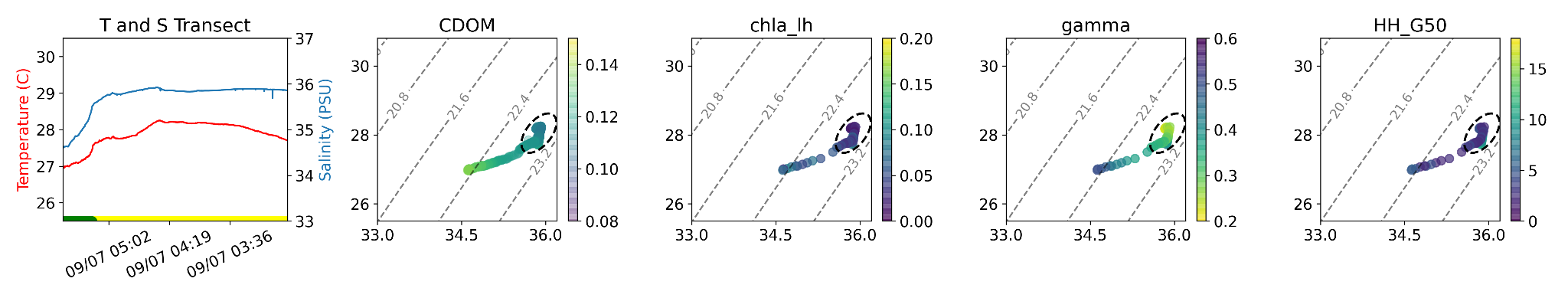

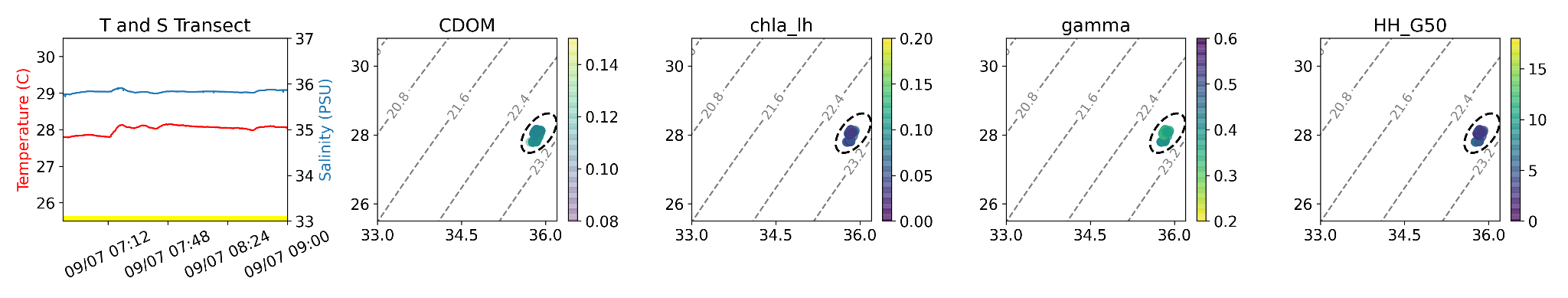

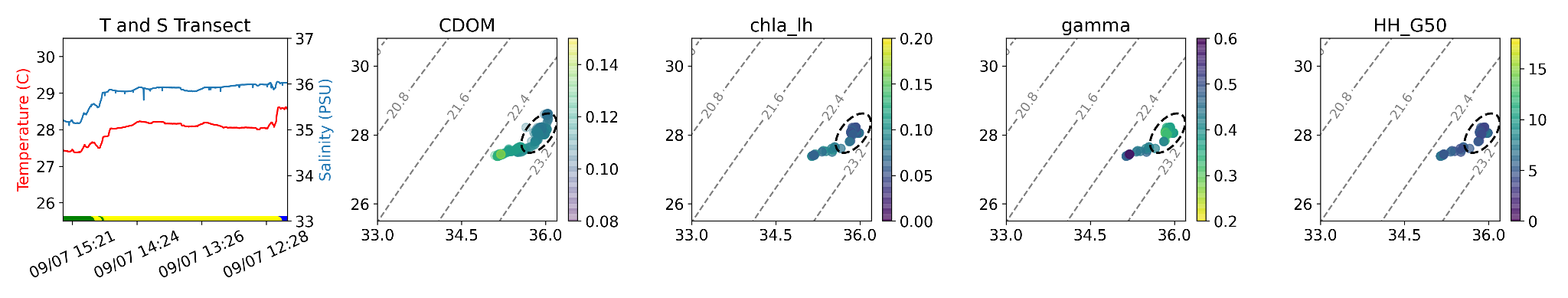

Figure S2. All transects shown along with optical parameters showing both transect values across time and plotted on a T-S diagram. Our first transect (morning to midday) transited from slope water into the Gulf Stream just before the eddy passed. Our second transect (midday to afternoon) was just as the eddy entered our study area and appeared to be just on the northern edge of the eddy, likely entirely in the warm streamer of the eddy with the highest temperatures of the cruise recorded here. Our third transect (sunset to midnight) and fourth (midnight to early morning) captured the beginning of the eddy, transiting from shelf water, through possibly slope water and into a mixture of eddy and Gulf Stream water. Transect five (morning to midday) made a clear transition from shelf water to eddy to Gulf Stream. Transect six (midday to afternoon) transits from shelf water into the eddy but passes another transition that appears to be the remains of the warm streamer that was pulled into the eddy. Transect seven (sunset to midnight) moves from shelf water into the eddy. Transect eight appears to be entirely in the eddy, and transect nine transits from the shelf water, through the middle of the eddy, and just into the Gulf Stream water.

Transect 1

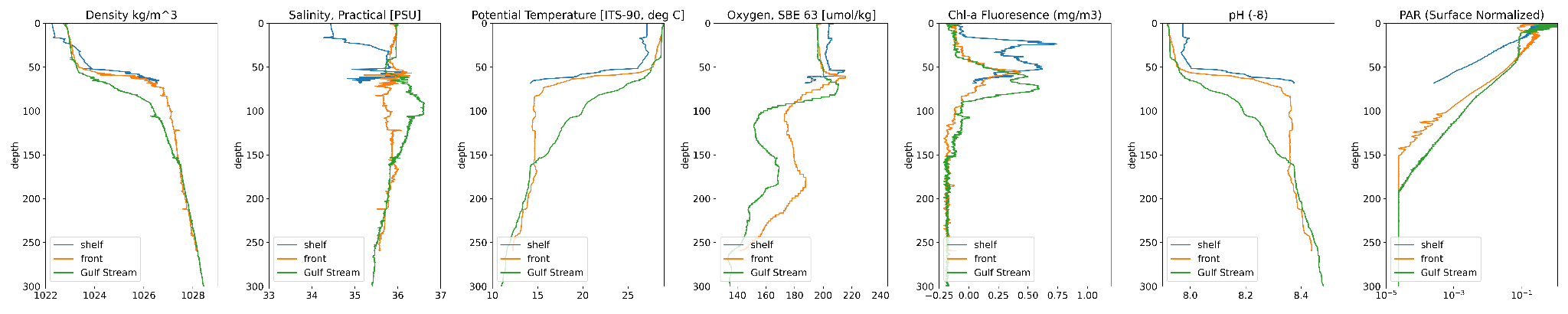

Transect 6

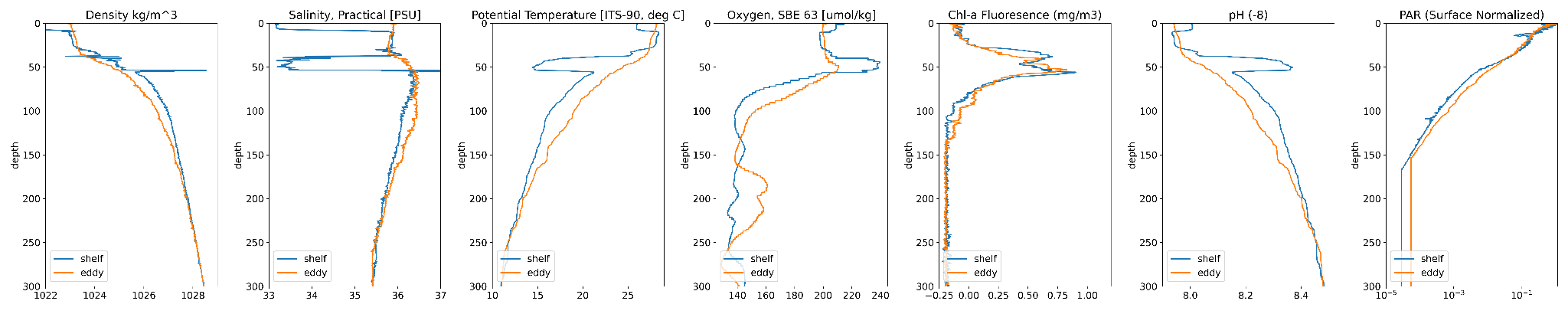

Transect 9

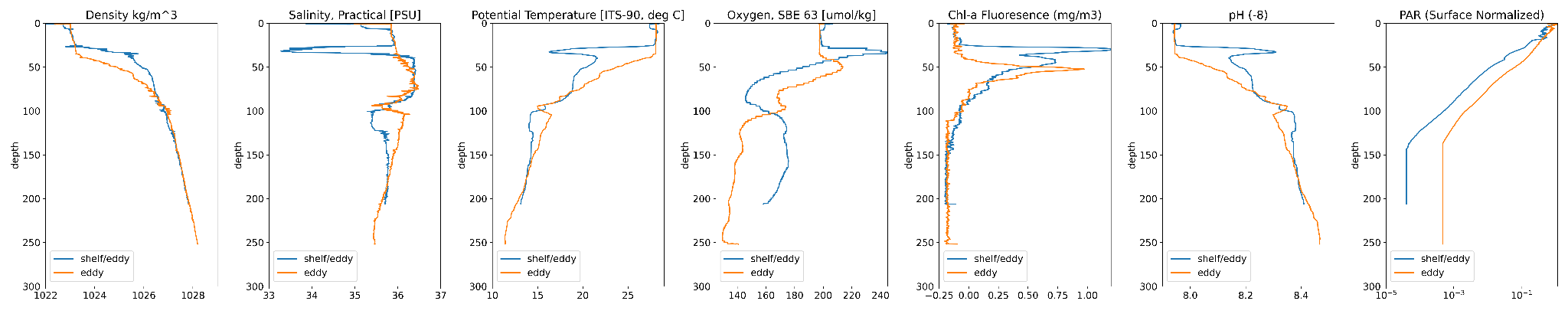

**Figure S3**. All CTD data from the cruise

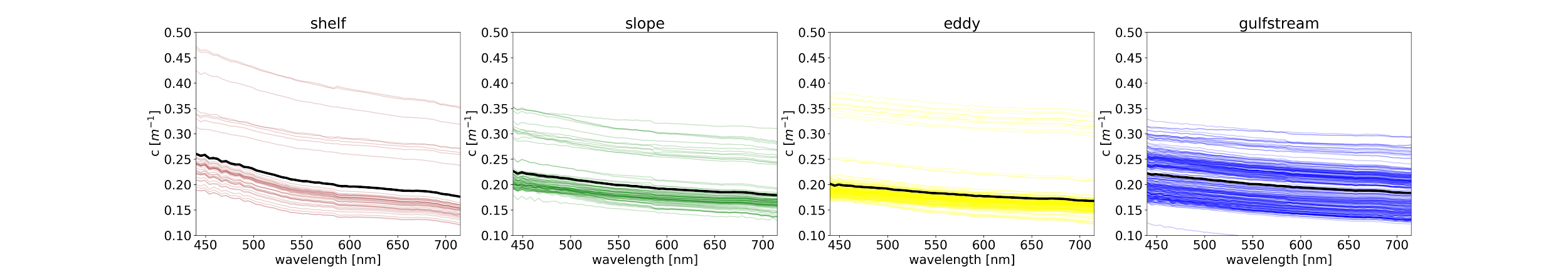

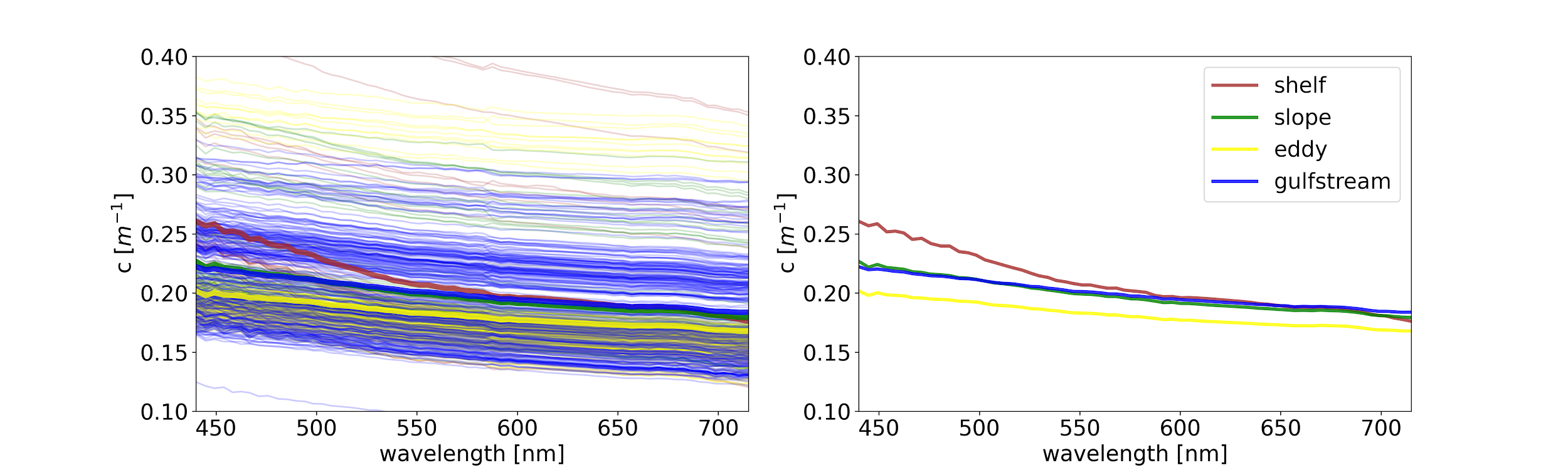

**Figure S4**. Attenuation spectra for all water masses.

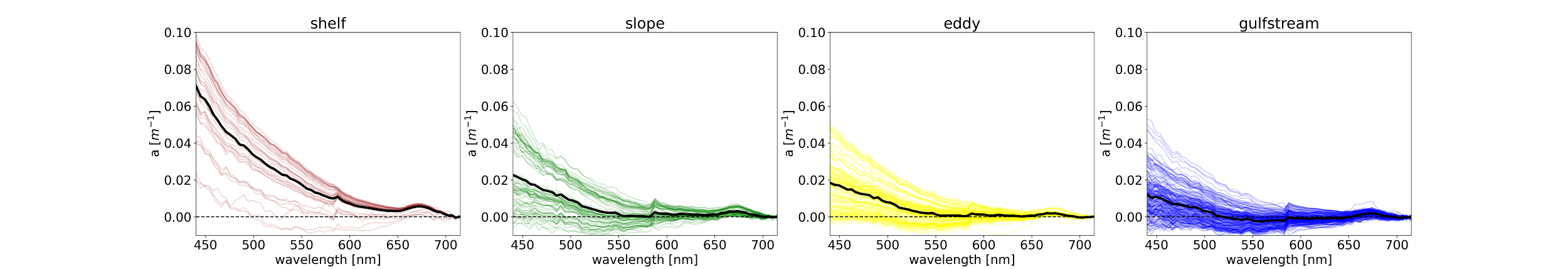

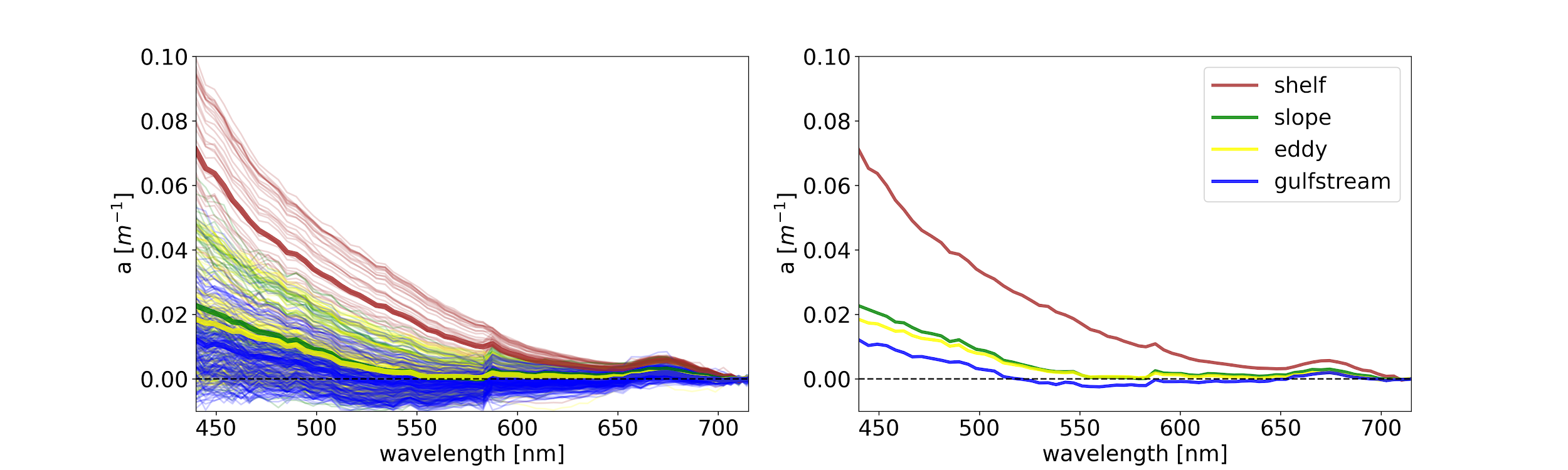

**Figure S5**. Absorption spectra for all water masses.

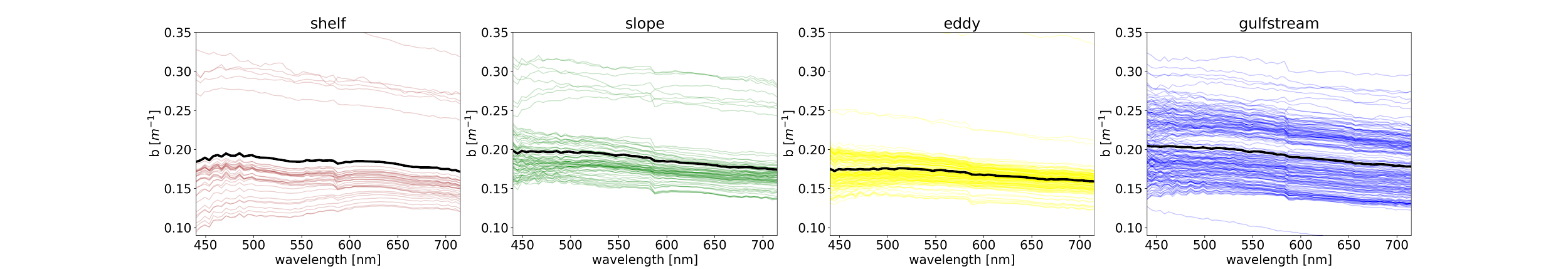

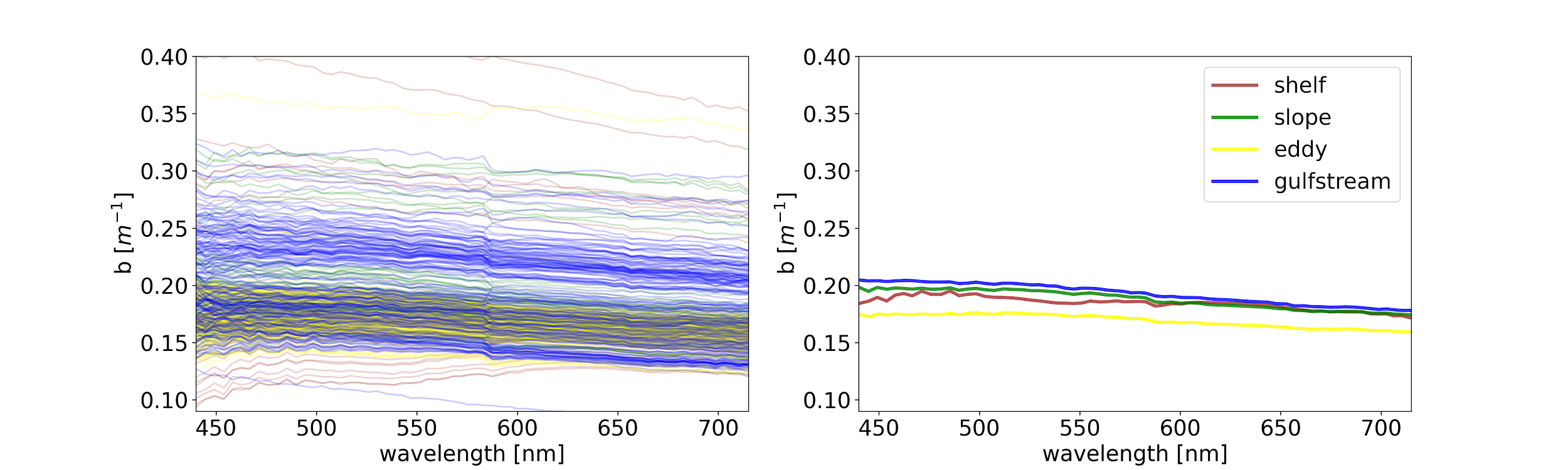

**Figure S6**. Scattering spectra for all water masses.

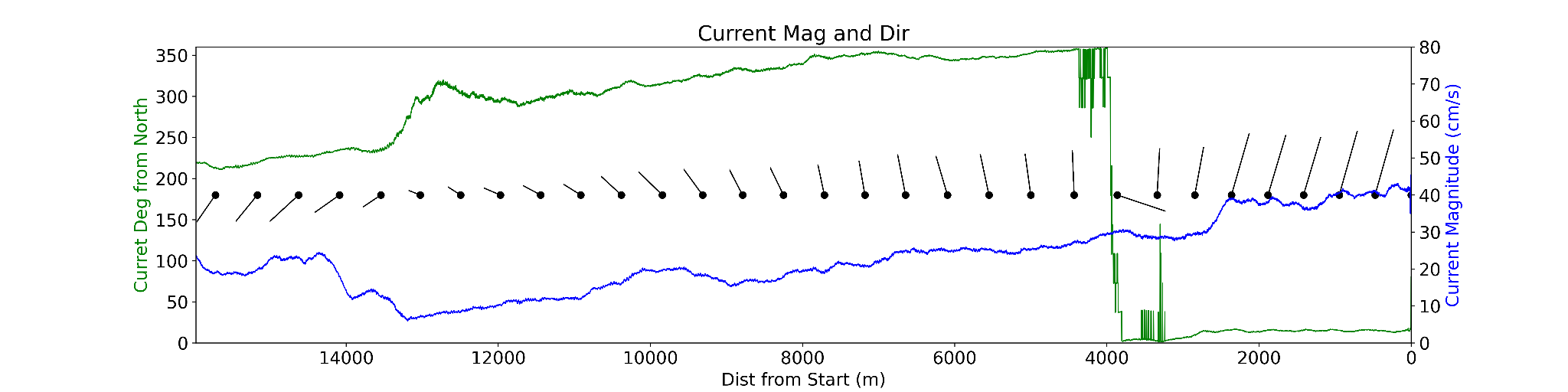

**Figure S7**. Magnitude and direction of the current on Transect 9 as measured by ADCP show a near complete reversal of the current indicating a slow (~25cm/s) continued cyclonic rotation. This panel shows current magnitude (blue) and direction (green) at a depth of 10 meters. Arrows show both direction and magnitude.
